# Repurposing the diatom periplastidial compartment for heterologous terpenoid production

**DOI:** 10.64898/2025.12.05.692597

**Authors:** Payal Patwari, Florian Pruckner, Luca Morelli, Michele Fabris

## Abstract

Diatoms are promising microorganisms to provide sustainable routes for photosynthetic terpenoid production from CO₂, yet their potential for compartmentalized engineering remains largely unexplored. Here, we systematically profiled the biosynthetic capacity of *Phaeodactylum tricornutum* by targeting representative synthases for hemi, mono, sesqui, and tetraterpenoids to the cytosol, chloroplast, and periplastidial compartment (PPC). This comprehensive analysis revealed that all major prenyl phosphate precursors, DMAPP, GPP, FPP, and GGPP, are accessible in all compartments, including in the PPC and can sustain heterologous flux without major physiological penalties, although production efficiency varies across compartments and product classes. By determining precursor availability, we propose the diatom PPC as a minimal engineerable organelle directly interfaced with a eukaryotic chloroplast. Moreover, we highlight its utility as a unique interface to investigate metabolic exchange between the MVA and MEP pathways. These findings provide a systematic framework for compartment-specific terpenoid engineering in diatoms and open new opportunities for modular pathway assembly and synthetic biology in photosynthetic eukaryotes.

## Introduction

Terpenoids are a vast and diverse class of metabolites with applications spanning pharmaceuticals, nutraceuticals, and industrial chemistry, representing a projected global market value exceeding 2B USD by 2030^1^. Direct sourcing of terpene-based products from plants and other native producers is often challenging, cost- and resource-ineffective and often raises sustainability concerns^2^. Heterologous production in engineered microorganisms is often considered a valid, scalable and sustainable alternative^3^.

The commercially viable production of heterologous terpenoids through metabolic engineering is still limited by several intrinsic challenges. These include interference and down-regulation of heterologous pathways by host regulatory networks^4,5^, restricted diffusion of precursors, intermediates, and enzymes within the complex cellular matrix^6^, strong competition from endogenous metabolic routes utilizing the same precursors, and robust homeostatic mechanisms that counteract the perturbations required for high-titre biosynthesis^7,8^. Furthermore, some isoprenoid precursors can become problematic when over-accumulated. For instance, farnesyl diphosphate (FPP) and related prenyl diphosphates are highly reactive and, if not efficiently channelled into downstream pathways, may interfere with protein prenylation, or trigger cytotoxic effects. Such imbalances can impose a metabolic burden on the host cell, as shown in studies where excess FPP accumulation reduced growth or viability^9,10^.

One strategy to address these challenges is to compartmentalize metabolite production into subcellular or synthetic spaces, separating heterologous pathways from competing host metabolism while maintaining resource access. In yeast, this has been achieved using prokaryotic nano compartments^11^ and artificial self-assembling organelles^12^, and by engineering existing organelles, such as peroxisomes^13^. Similar concepts have also been applied with encapsulins and bacterial microcompartments in other hosts^14,15^.

Diatoms are photosynthetic microalgae ubiquitously distributed in marine and freshwater habitats and have been investigated as novel platforms for terpenoid engineering, demonstrating promising proof-of-concepts for the heterologous production of monoterpenoids^16,17^, sesquiterpenoids^18^, triterpenoids^19,20^ and cannabinoids^21^ autotrophically from CO_2_, with inexpensive growth requirements, such as seawater and light.

Unlike plants, yeasts, prokaryotes, green algae and other established production hosts, diatoms offer unique opportunities for compartmentalized biosynthesis. Because they have originated through a secondary endosymbiotic event^22^, diatoms possess the periplastidial compartment (PPC). This consists in an additional intracellular space which surrounds the chloroplast, representing the remnant cytosol of an endosymbiont red alga that underwent progressive gene transfer and metabolic reduction^23,24^. Presently, only a small number of proteins are thought to have persisted in the PPC^25^, suggesting that limited metabolic activity might occur in the PPC, potentially involving heme biosynthesis, carbon concentration mechanisms, or lipid transfer^25–27^. Thus, the diatom PPC can be defined as a minimal eukaryotic cytoplasm tightly connected with the chloroplast, in a fast-growing photosynthetic cell. In a synthetic biology perspective, this organelle is an uncharted subcellular environment and an ideal candidate for compartmentalized, light-driven biosynthesis.

The potential to exploit the PPC is further enhanced by diatoms’ distinctive terpenoid metabolism, which operates through mechanisms fundamentally different from conventional production platforms. Peculiarities include the presence of novel and unusual enzymes^28,29^ and precursors^16^, which set them apart from conventional production platforms and can enable novel engineering strategies^30^. In most of the diatoms species characterized to date, the 5-carbon isoprenoid precursors isopentenyl diphosphate (IPP) and dimethylallyl diphosphate (DMAPP) are synthesized via the mevalonate (MVA) pathway in the cytosol, with some steps also in the endoplasmic reticulum (ER)^20^ and possibly other organelles^31^, and the methyl-erythritol 4-phosphate (MEP) pathway in the plastid^32^. IPP and DMAPP can be condensed by prenyl transferases to form 10-, 15-, and 20-carbon prenyl phosphates, geranyl diphosphate (GPP), farnesyl diphosphate (FPP), and geranylgeranyl diphosphate (GGPP) respectively^33–35^.

It has been observed that in several organisms which harbour both pathways, these interact and exchange metabolites. A metabolic crosstalk occurring between the MVA and the MEP pathway, has been shown to operate in plants^36,37^. However, decades of research have not yet elucidated the elusive molecular mechanisms underpinning this crosstalk, which can hold the key to accurately manipulate their regulation and thus their metabolic flux. In diatoms, the PPC surrounds the chloroplast^38^, and we hypothesize that, if exchanges occur, this necessarily mediates the interaction between MEP and MVA pathway. In this scenario, the diatom PPC provides a unique opportunity to investigate this exchange of isoprenoids between the chloroplast and other compartments, and to exploit this mechanism to directly channel these precursors in heterologous pathways built ex-novo in the PPC.

In this study, we systematically investigated whether the PPC can support the biosynthesis of heterologous terpenoids in *P. tricornutum*, starting from each prenyl phosphate precursor DMAPP, GPP, FPP and GGPP. We chose isoprene, geraniol, zizaene, and phytoene as model products with commercial applications, spanning four of the main classes of terpenoids, and the main classes of prenyl phosphate precursors. Isoprene is a volatile hemiterpene widely used in the production of rubber, adhesives and has potential in bio-based fuel production^39–42^. Geraniol, a volatile monoterpenoid alcohol with a rose-like fragrance^43^, is commonly used as flavors and fragrances^44^ and pharmaceuticals, serving as a precursor of monoterpene indole alkaloids for anti-cancer drugs^45^ and has exhibited anti-inflammatory and antimicrobial properties^46^. Zizaene is a volatile sesquiterpene and a biosynthetic precursor to khusimol^47^, a key fragrance ingredient in perfume industry and its derivatives are also investigated for antimicrobial and anti-inflammatory activities^49,50^. Phytoene is a non-volatile tetraterpene and an early precursor of carotenoids such as lutein, zeaxanthin and β-carotene^51^, used as natural antioxidants in nutraceuticals and cosmetics^52^. These compounds were used as model heterologous products and proxies to infer the availability of their specific precursors in the cytosol, chloroplast and PPC. Through this, we aim to determine whether the PPC can be repurposed as a productive platform for high value terpenoid synthesis.

## Materials and Methods

### Strain, cultivation and growth conditions

*P. tricornutum* strain CCMP632 was obtained from the National Center for Marine Algae and Microbiota (Bigelow Laboratory for Ocean Sciences, USA) and sub-cultured once a week in Enriched Seawater Artificial Water (ESAW) medium, supplemented with 100 µg/mL Zeocin (Lifetech, USA) for transgenic lines. Solid medium plates consisting of half-strength ESAW with 1% agar and supplemented with 100 µg/mL Zeocin were used for selection, while liquid ESAW supplemented with 100 µg/mL Zeocin was used to grow all experimental cultures. In all experiments, cultures were grown in a shaking incubator (Innova S44i, Eppendorf, Germany) maintained at 21°C, under continuous light at 80 μmol photons m⁻² s⁻¹ and shaking at 95 rpm.

### Constructs assembly and generation of transgenic *P. tricornutum* cell lines

All episomes were assembled using the uLoop system, as previously described^53^. Full length DNA sequences encoding an isoprene synthase (EC: 4.2.3.27) from *Populus alba* fused with green fluorescent protein (ISPS-GFP, gene accession number: ABV04402.1) truncated of the plant targeting sequence^38^, a full length geraniol synthase (EC: 3.1.7.11) from *Catharanthus roseus* (GES, gene accession number: JN882024.1)^16^, a cytosolic zizaene synthase (EC: 4.2.3.-) from *Chrysopogon zizanioides* (ZS, gene accession number: HI931360.1)^48^, and a bacterial phytoene synthase (EC: 2.5.1.32) from *Pantoea ananatis* (CrtB, gene accession number: ADD79329) were codon-optimized for *P. tricornutum* using Benchling [Biology Software] (2025) and synthesized by Genewiz (USA, 2025). These genes were cloned into the pL0-R vector as “CD” parts and the sequences are listed in Table S3.

The pCAO-2 vectors carrying genes for cytosolic expression were assembled by combining the constitutive (*Phatr3_J49202)* or the inducible promoter (*pPhAP1, Phatr3_J49678*) promoter region^55^ (“AC”), the codon-optimized terpenoid gene (“CD”), the “DE” part (typically mVenus-stop codon), and the *FcpB* terminator (“EF”). An exception was ISPS-GFP, which included only the stop codon as the “DE” part, due to the presence of GFP within its coding sequence. To assemble episomes carrying genes to be targeted to either the PPC or the chloroplast, dedicated “CD” parts for terpene synthase coding sequences were assembled with a transit peptide sequence fused upstream of the start codon synthesized by Genewiz (USA). For targeting the terpene synthases to the PPC and chloroplast, 55 amino acids N-terminal transit peptide sequence of *Hsp70* (*Phatr3_J21519*) and 100 amino acids N-terminal sequence of the *γ* subunit of the diatom plastidial ATPase (*AtpC*, *Phatr3_J20657*) were used^56–58^ (Table S3). The pCAO-1 vectors were obtained from uLoop kit developers^59^ and included L1-1 PTCv2, containing a sh-*bleR* selection cassette, CEN/ARS/HIS region for episomal maintenance in diatoms and oriT. pCAO-3 and pCAO-4 carried spacer sequences. All four pCAO-1, -2, -3, and -4 were assembled to form the final L2 episomes, pCAe-1, which were subsequently conjugated into *P. tricornutum*. All uLoop parts used are described in Table S1.

Episomes were introduced into *P. tricornutum* using a bacterial conjugation approach adapted from^60^. A 50 mL culture was inoculated with exponentially growing *P. tricornutum* cells at an initial OD_750_ of approximately 0.03 and cultivated until reaching an OD_750_ of 0.3. For conjugation, six-well plates were prepared by adding 3 mL of a ½ ESAW medium supplemented with 5% LB and 1% agar to each well.

On the day before conjugation, cells in mid- to late-exponential phase (cell density of 1.0 × 10^8^ cells□m/L) were harvested by centrifugation at 4000 × g. The pellet from each 50 mL culture was resuspended in 500 μL of ESAW to adjust the cell concentration to 5 × 10^8^ cells□m/L, and 50 μL of this suspension was spotted onto the ½ strength ESAW agar six-well plates. The plates were dried and incubated under constant illumination of 80 μmol photons m⁻² s⁻¹ at 21°C overnight. The same day an overnight culture of *E. coli* Epi300 harboring both the conjugative plasmid pTA-MOB and the episomal cargo plasmid was prepared by shaking at 180 rpm at 37°C. This culture was then diluted 1:50 into 20 mL of fresh LB containing 20 μg m/L gentamicin and 50 μg m/L spectinomycin and grown to an OD_600_ between 0.8 and 1.0. The bacterial cells were pelleted by centrifugation (10 min at 3000 × *g*), the supernatant was discarded, and the pellet was resuspended in 250 μL of SOC medium. The next day a 50 μL aliquot of the resuspended *E. coli* culture was added directly onto each dried diatom spot and left to dry under sterile conditions. Plates were first incubated in the dark at 30°C for 90 minutes, then transferred to a shaking incubator (95 rpm), under continuous light (80 μmol photons m⁻² s⁻¹) at 21°C for recovery of three days.

Next, 1 mL of ESAW medium was added to each well to resuspend the cells, and 400 μL was transferred to selective ½ ESAW agar plates containing 100 μg/mL Zeocin and 50 μg/mL kanamycin to suppress bacterial growth. Transformed *P. tricornutum* colonies were visible after 2–3 weeks. Each colony was resuspended in 200 μL of ESAW containing 100 μg/mL Zeocin in 96-well plates (Nunclon-treated; Thermo Fisher Scientific, USA). After a week the plates were subcultures with 20 μL of culture and 180 μL ESAW containing 100 μg/mL Zeocin and after further 4 days the transformants were analyzed using a Guava easyCyte HT flow cytometer (CyTek Biosciences, USA). Fluorescent proteins were excited with a 488 nm laser, and FITC emission was detected in the green channel (525/40 nm filter). Each sample was analysed in triplicates, recording 5,000 events per well at a flow rate of 0.59 μL/s. Three to four colonies exhibiting the highest signal were selected for experiments.

### Confocal laser microscopy

Cells expressing heterologous terpene synthases were prepared at a cell density of 1.0 × 10^5^ cells□m/L for live imaging. Imaging was performed using a Nikon A1 confocal microscope (Nikon, Japan) equipped with a Plan Apo λ 100× oil-immersion objective lens (numerical aperture 1.49). Images were captured at a resolution of 1024×1024 pixels. The mVenus fluorophore was excited using a 488 nm laser, while chlorophyll autofluorescence was excited with a 561 nm laser. Emission signals were collected in the 500–520 nm range for mVenus and 625–720 nm for chlorophyll. Image processing and analysis were carried out using ImageJ software.

### Isoprene extraction and detection by GC-MS and quantification by GC-FID

Isoprene-producing transgenic diatoms were first grown to preculture containing 50 µg/mL Zeocin. Cell density was determined using flow cytometry, and cultures were then resuspended in fresh ESAW medium supplemented with 15□µM H₂CO₃. The cell density was adjusted to 10,000 cells/µL, and 5.9□mL Exetainer vials (Labco, USA) were inoculated with the resuspended cultures. Cultures were grown for 3 days under constant light (24:0 light:dark), at 21□°C, 95□rpm, and 80 μmol photons m⁻² s⁻¹. After the incubation period, vials were incubated at 45□°C for 20 minutes with shaking at 85□rpm to enrich isoprene in the vial headspace. Immediately after heating, 1.5□mL of the headspace gas was manually sampled using a 2.5□mL gastight syringe (Avantor, USA) and injected into a gas chromatograph for analysis. Gas chromatographic analysis was performed on an Agilent 7890B GC system equipped with an Agilent J&W DB-5 column (Agilent, USA) and a flame ionization detector (FID). The injector temperature was set to 280°C with an inlet pressure of 11.084□psi. A pulsed split injection was used with an injection pulse pressure of 30□psi for 0.4□min, a split ratio of 25:1, and a split flow rate of 25□mL/min. A septum purge flow of 3□mL/min was maintained. The oven temperature program was as follows: initial temperature of 30°C, ramped to 34°C at 2°C/min, followed by a rapid increase to 45°C at 110°C/min, which was held for 1 minute. The nitrogen carrier gas was maintained at a flow rate of 1□mL/min under 11.084□psi. Detector conditions were FID temperature at 365°C, airflow at 300□mL/min, hydrogen fuel flow at 30□mL/min, and nitrogen makeup flow at 20□mL/min.

### Geraniol and zizaene extraction and detection by GC-MS and quantification by GC-FID

To capture geraniol and zizaene, 2 mL isopropyl myristate (IM) was added to 50 mL diatom cultures at the time of inoculation. After 12 days of incubation period, the IM overlay was harvested by centrifugation at 4,500 × *g* for 10 minutes. The collected IM phase was then subjected to gas chromatography–mass spectrometry coupled with a flame ion detection (GC–MS/FID) (Agilent, USA) for compound analysis. Helium served as the carrier gas at a constant flow rate of 1.0 mL/min, and injection volume of 1 μL. The chromatographic separation was done with a nonpolar HP-5 column (5%-phenyl)-methylpolysiloxane (30.0 m × 0.32 mm × 0.25 μm). The oven temperature was initially held at 35°C for 3 minutes, then ramped at 20°C/min to 320°C and maintained for an additional 5 minutes. The mass spectrometer operated in electron impact (EI) mode at 70 eV. The injector and ion source temperatures were set to 280°C and 230°C, respectively. Geraniol and α-cedrene were quantified in Selective Ion Monitoring (SIM) mode, targeting the ion at *m/z* 69.1 for geraniol and *m/z* 119 for α-cedrene with a dwell time of 20 ms per ion. Metabolite concentrations were determined by comparing peak areas to a standard calibration curve generated using known concentrations of geraniol and α-cedrene (Sigma-Aldrich, USA), closely related to zizaene, as a proxy for zizaene, since the authentic standard of the latter is not commercially available.

### Phytoene extraction and quantification by HPLC

Phytoene was extracted and quantified from freeze-dried algal pellet using a biphasic solvent system. Samples were placed in 2 mL microcentrifuge tubes with 400□µL of methanol (MeOH). After vortexing for 10□s, samples were additionally disrupted by vigorous shaking on an automated rocking system for 10□min at 4°C. Subsequently, 400□µL of Tris-HCl buffer (100 mM, pH 7.5) were added, and the mixture was homogenized again under the same conditions. Next, 800□µL of chloroform were added, followed by a third round of homogenization. The samples were then centrifuged at 4°C for 5□min at maximum speed, and the lower organic phase was collected and evaporated under nitrogen flux

Dried extracts were reconstituted in 200 µL of methanol:acetone (80:20) prior to analysis by HPLC. Pigment separation was performed on a YMC column (150 x 3.0 mm, S-3µm) column using a binary gradient system with Solvent A: methanol/water (80:20, v/v) containing 0.2% (w/v) ammonium acetate, and Solvent B: tert-methyl butyl ether (MTBE). The elution profile was as follows: 0–3 min, 90% A; 3–10 min, linear gradient to 50% A; 10–22 min, linear gradient to 5% A; 22–25 min, held at 5% A; 25–27 min, returned to 90% A; 27–30 min, re-equilibration at 90% A. Detection was carried out with a photodiode array detector monitoring absorbance at 286 nm.

### Photosynthetic measurements

On the final day of each experiment, photosynthetic activity of the transgenic lines was assessed using an AquaPen-C fluorometer (Photon Systems Instruments, Czech Republic). Measurements were performed on light-adapted samples to determine the effective quantum yield of photosystem II (ΦPSII). This parameter was calculated as the difference between the maximum fluorescence yield after a saturating light pulse (Fm′) and the steady-state fluorescence (F), divided by the maximum fluorescence yield (Fm′). In this context, F represents the fluorescence emitted by the sample under continuous actinic light, while Fm′ corresponds to the peak fluorescence recorded immediately after the application of a saturating pulse of light.

## Results and discussion

### Heterologous terpene synthases can be targeted to the cytosol, chloroplast and PPC

We systematically targeted four heterologous terpene synthases that specifically use the entire range of prenyl phosphate substrates (DMAPP, GPP, FPP and GGPP) to the cytosol, chloroplast and PPC of *P. tricornutum*. Specifically, we expressed *P. alba* isoprene synthase (ISPS), *C. roseus* geraniol synthase (GES), *C. zizanioides* zizaene synthase (ZS) and *P. ananatis* phytoene synthase (crtB). Heterologous enzymes targeted to the cytosol, were expressed without transit peptides, whereas to favour their import in the chloroplast and PPC, we fused the bipartite transit peptide of the gamma subunit of the thylakoid ATP synthase (AtpC, Phatr3_J20657)^58^ or the bipartite transit peptide of the PPC-located Heat shock protein 70 (HSP70, Phatr3_J21519)^56,57,62^ to their N-terminal portion, respectively. To track their subcellular localization, we fused a yellow fluorescent protein (mVenus) to the C-terminus of GES, ZS and crtB, while a green fluorescent protein (GFP) was embedded in ISPS coding sequence^54^. As controls, we expressed mVenus either free or fused to AtpC or HSP70 transit peptides (Fig. S1). Since phytoene is endogenously produced from GGPP in the chloroplast of *P. tricornutum*^63^, and rapidly converted into downstream carotenoids, we decided to express crtB in the cytosol and in the PPC, but not in the chloroplast.

When introduced in *P. tricornutum*, all four terpene synthases were successfully expressed and accumulated in their designated subcellular space (Fig. 2). Those lacking diatom transit peptides accumulated in the cytosol as shown by an even distribution of mesh-like green fluorescence pattern throughout the cytoplasm^62^. GES-mVenus, which still contains the plant plastid targeting sequence at its N terminal, which is not recognized in *P. tricornutum*^16,64^, as the fusion protein was directed into the cytoplasm. We observed a green fluorescence pattern resembling the “blob-like” structure typical of fusion fluorescent proteins containing the HSP70 target peptide^43^. Recombinant enzymes fused with to AtpC target peptide showed distinct green fluorescence overlapping with chlorophyll autofluorescence in the chloroplast stroma, indicating plastidial localization^58,62^ (Fig. 2).

**Figure 1.**
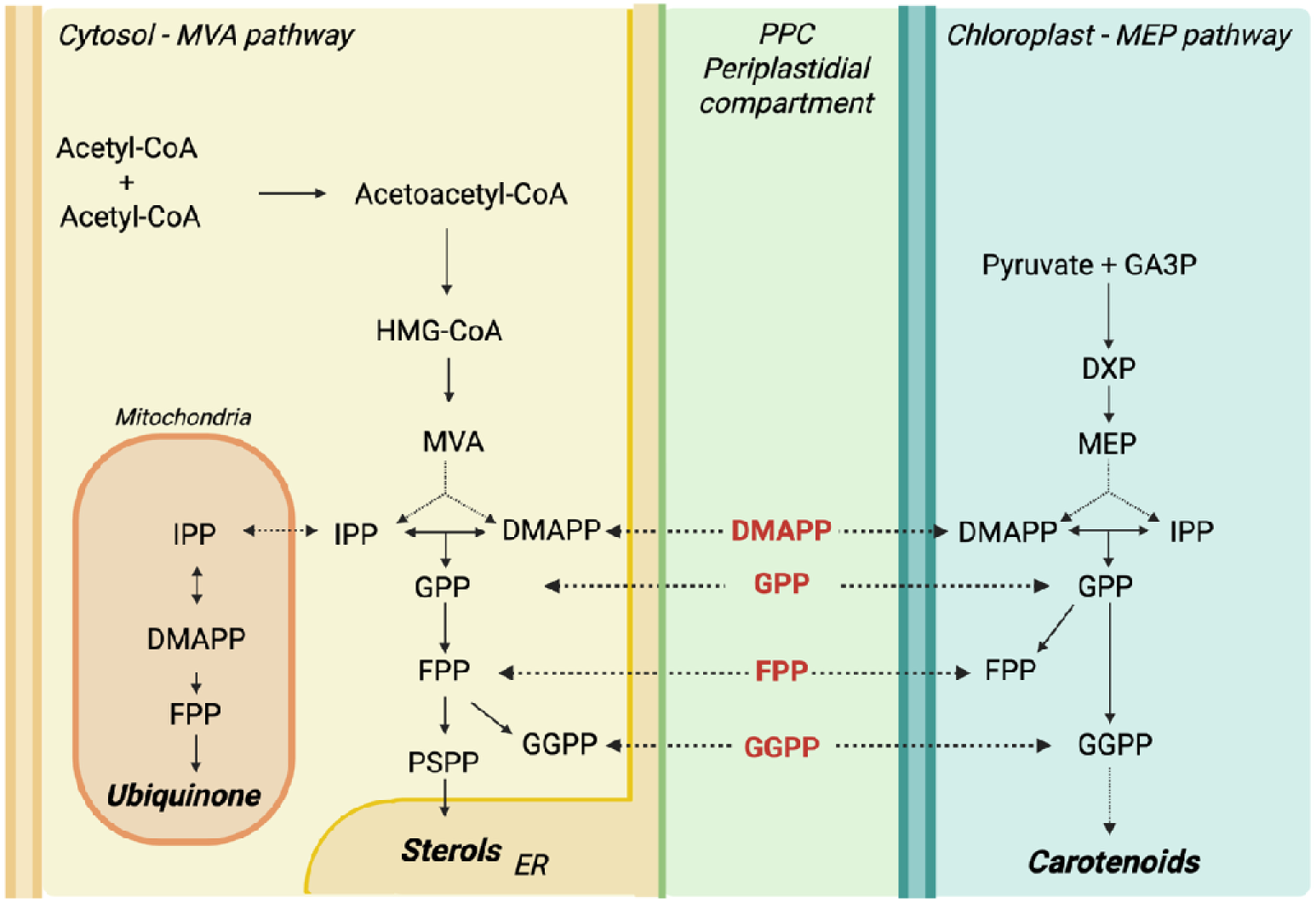
Schematic overview of the terpenoid metabolic network in *P. tricornutum* and its associated membrane architecture. The methylerythritol 4-phosphate (MEP) pathway is localized in the chloroplast, which is enclosed by the periplastidial compartment (PPC), surrounded by the endoplasmic reticulum (ER). The mevalonate (MVA) pathway operates across the cytosol, ER and potentially other organelles, but for simplicity here is represented in the cytosol. Both the MEP and MVA pathways produce the prenyl phosphates dimethylallyl diphosphate (DMAPP) and isopentenyl diphosphate (IPP), which serve as precursors for geranyl diphosphate (GPP), farnesyl diphosphate (FPP), and geranylgeranyl diphosphate (GGPP). FPP from the cytosol is directed toward sterol biosynthesis in the ER, while carotenoid biosynthesis occurs within the chloroplast from GGPP. Dashed lines indicate hypothetical exchange of prenyl phosphate precursors. Created in BioRender. Morelli, L. (2026) https://BioRender.com/o88epb9

**Figure 2:**
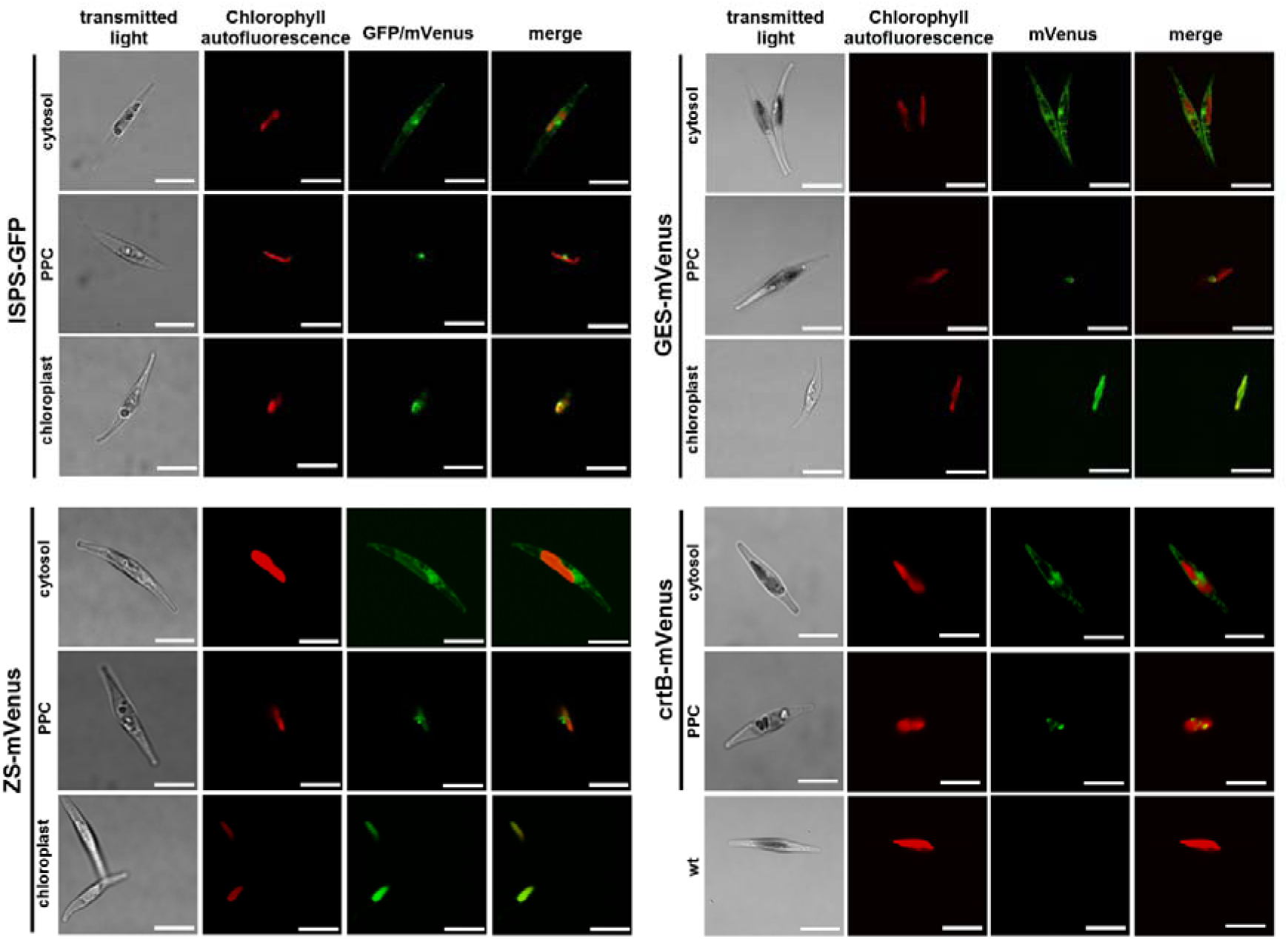
Subcellular localization of heterologous terpene synthases in *P. tricornutum*. Representative confocal images of diatom cell lines expressing the fusion terpene synthases ISPS_GFP, GES_mVenus, ZS_mVenus and crtB_mVenus (targeted to cytosol); AtpC_ISPS_GFP, AtpC_GES_mVenus, AtpC_ZS_mVenus (targeted to chloroplast); Hsp70_ISPS_GFP, Hsp70_GES_mVenus, Hsp70_ZS_mVenus and Hsp70_crtB_mVenus (targeted to PPC) and wild type (wt). Each image is representative of at least three independent cell lines per construct. Scale bars represent 5 µm.

### Prenyl phosphate pools are available in the cytosol, chloroplast and PPC and can be used to synthesize heterologous terpenes

While isoprenoids biosynthesis is known to occur in the chloroplast, cytosol and ER, no enzymes involved in isoprenoids biosynthesis has been reported nor have been computationally predicted^61^ to localize in the PPC of diatoms. Prenyl transferases are the enzymes responsible to convert IPP and DMAPP into longer-chain intermediates such as GPP, FPP, and GGPP. In *P. tricornutum*, five genes have been annotated as putatively encoding prenyl transferases (*Phatr3_J19000, Phatr3_J15180, Phatr3_J16615, Phatr3_J47271, and Phatr3_J49325*). To date, their subcellular localization has only been determined from computational predictions, none of which place them in the PPC^41^ and their potential functional contribution to PPC metabolism remains unresolved.

In our experimental setup, each terpene synthase was expressed under the control of identical promoters and terminators, differing only in their transit pre-sequence. Exploiting the higher consistency in expression levels allowed by extrachromosomal episomal expression compared to random chromosomal integration^17^, this design enables direct comparisons of enzyme activity across compartments. Terpenoid production and fluorescent protein signals were interpreted as semiquantitative indicators of local precursor availability and TPS accumulation, respectively. However, this comparison does not account for intrinsic compartment-specific factors such as pH^65,66^, redox state, autofluorescence, and fluorophore maturation, that may influence fluorescence, nor for differences in metabolic context that could affect enzymatic conversion efficiency.

#### DMAPP availability and isoprene production via heterologous ISPS activity

Isoprene is formed from DMAPP catalyzed by ISPS enzymes, although it can also be synthesized in absence of canonical ISPSs^67^. The genome of *P. tricornutum* does not encode conventional isoprene synthase enzymes^16^, although small amounts of isoprene production have been reported in this species in certain conditions^68^. Diatoms strains expressing ISPS-GFP targeted to the cytosol, PPC, and plastid in gas-tight sealed vials for four days, all produced isoprene, which accumulated in the headspace of the vial. Isoprene production was confirmed by GC-MS (Fig. S2) and quantified by GC-FID (Fig. 3A). Cultures expressing an ISPS-GFP in the chloroplast emitted isoprene with a yield of 0.708 mg/L, significantly higher than strains expressing the same fusion enzyme in the cytosol (0.1 mg/L) and PPC (0.06 mg /L). In contrast to the engineered *C. reinhardtii* strains that reached up to 334 mg/L isoprene under optimized CO_2_-fed photobioreactor conditions, the yields we recorded remain substantially lower. These differences likely reflect limited DMAPP precursor supply in *P. tricornutum*, since no upstream pathway enzymes were overexpressed to boost precursor flux.. Control strains expressing a cytosolic mVenus marker emitted only traces amount of isoprene, consistent with previous reports^42^, confirming that large part of the isoprene emission observed in the experimental cell lines originated from ISPS-GFP activity. The rates of isoprene emission were proportional to the abundance of ISPS-GFP in each compartment, as quantified by median GFP fluorescence at the time of harvest, which was surprisingly higher in the chloroplast (Fig. 3A). These findings indicate that DMAPP pools are accessible in the cytosol, PPC, and chloroplast, that ISPS-GFP is functionally active in all three subcellular locations, and that isoprene production is directly proportional to the GFP fluorescent signal detected at harvest.

**Figure 3:**
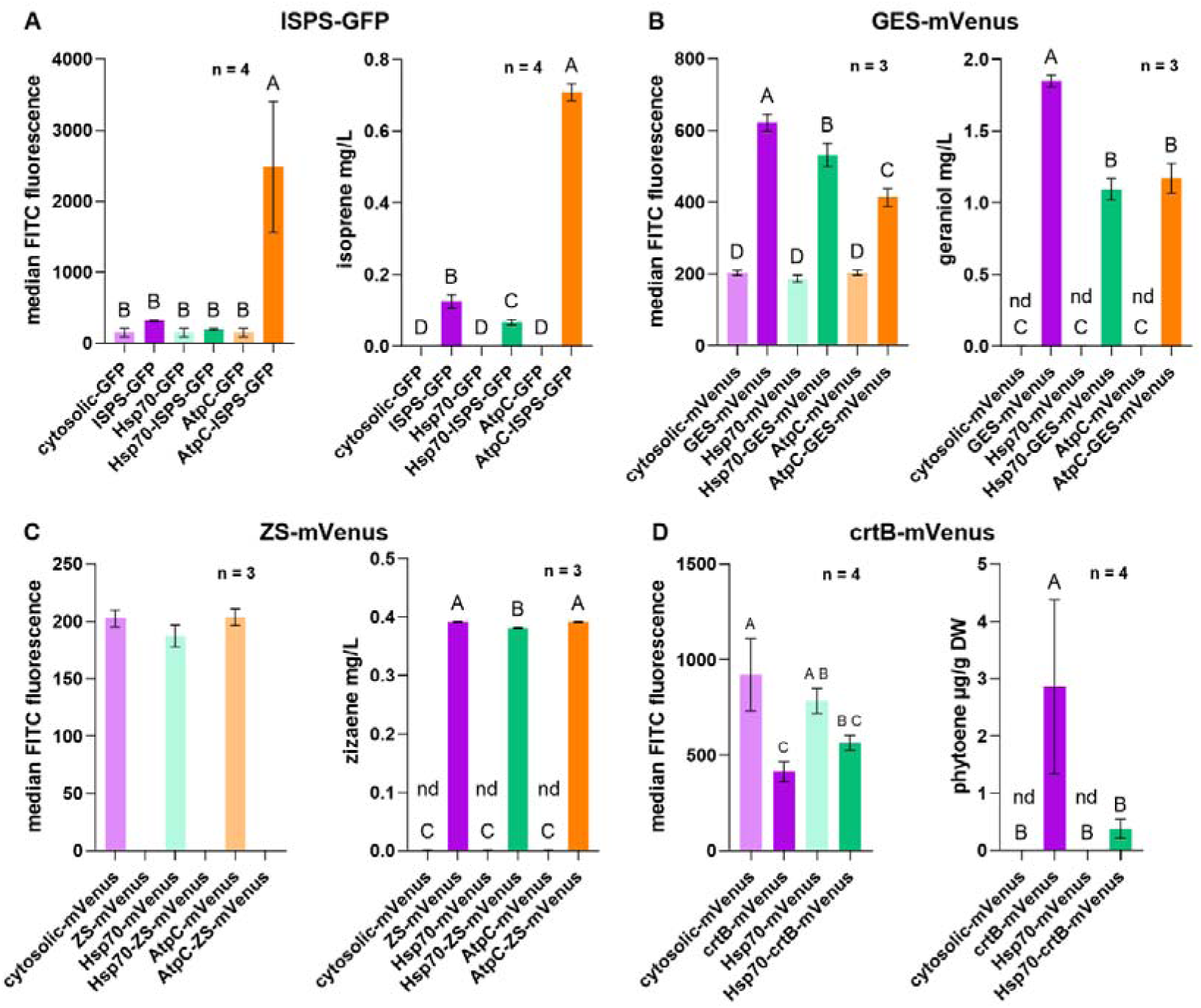
Heterologous terpene synthases expression levels and production of different classes of terpenoids in the cytosol, PPC, and chloroplast. Each panel represents (left) quantification of median GFP or mVenus fluorescence intensity normalized to cell density, measured by flow cytometry (5000 events) on the last day of the experiment, and indicative of the amount of terpene synthases fused to fluorescent proteins in each subcellular location; (right) heterologous metabolite quantification. **(A)** Isoprene production (ISPS-GFP); **(B)** Geraniol production (GES-mVenus) **(C)** Zizaene production (ZS-mVenus) **(D)** Phytoene production (crtB-mVenus). Statistical differences between groups were determined by one-way ANOVA, with different letters indicating significant differences (*p*-value < 0.05); nd = not detected.

The availability of DMAPP in the cytosol and chloroplast of *P. tricornutum* was expected, since the diatom possesses two distinct enzymes responsible for the biosynthesis of DMAPP and its isomer IPP. An hydroxymethylbutenyl diphosphate reductase (HDR, EC: 1.17.1.2, Phatr3_J41845) converts (E)-4-hydroxy-3-methylbut-2-enyl diphosphate into IPP and DMAPP, has a high-confidence chloroplast transit peptide^16^, and is part of the plastidial MEP pathway. Contributing to the cytosolic DMAPP moiety, a diphosphomevalonate decarboxylase (MDPC, EC: 4.1.1.33, Phatr3_EG02359), part of the MVA pathway, converts 5-diphosphomevalonate into IPP and does not include any evident transit peptides^16^. A multifunctional isopentenyl diphosphate isomerase/squalene synthase (EC: 5.3.3.2/2.5.1.21, PtIDISQS, Phatr3_EG02290) without predicted transit peptides catalyzes the isomerization of cytosolic/ER IPP to DMAPP^29^, while a plastidial IDI enzyme (EC: 5.3.3.2, Phatr3_J12533) carries out the isomerization between IPP and DMAPP derived from the MEP pathway. While the subcellular localization of these enzymes has not been experimentally reported, their predicted organization, based on sequence analyses, resembles that of plants, except for the multifunctional enzyme PtIDISQS. Hence, the presence of distinct pools of DMAPP/IPP was expected in the cytosol and chloroplast, while the presence of DMAPP in the PPC, in amounts sufficient to sustain isoprene synthesis was surprising, and suggests the existence of a metabolite exchange mechanism between chloroplast and cytosol, or of an unknown enzymatic source of DMAPP in the PPC.

#### GPP availability and geraniol production via heterologous GES activity

While diatoms have been reported to produce some monoterpenes naturally^70,71^, no monoterpene synthases are encoded in the genome of *P. tricornutum* ^16^. This species and the strain (Pt1) investigated here, do not synthesize endogenous monoterpenoids in the laboratory conditions used in this and other works^16,72^. We previously demonstrated that the expression of a cytosolic GES-mVenus fusion enzyme enables the production of heterologous geraniol from endogenous cytosolic GPP pools^16,72^. When expressed in the cytosol, chloroplast and PPC, GES-mVenus was functional in all cell lines, as these produced geraniol in all three subcellular compartments (Figs. 3B, S3). After 13 days of cultivation, the highest titres of geraniol were achieved in cell lines expressing GES-mVenus in the cytosol (1.848 ± 40.77 mg/L), followed by cell lines expressing GES-mVenus in the chloroplast (1.170 ± 102.40 mg/L) and in the PPC (1.094 ± 75.20 mg/L) (Fig. 3B). The relative yields obtained were proportional to the GES-mVenus signal detected at harvest, which were highest in cytosol and lowest in the chloroplast (Fig. 3B), suggesting the production correlated primarily with enzyme abundance rather than with precursor limitation as also observed for isoprene (Fig.3A) and in other studies^15,16,62^. Yields from cell lines expressing GES-mVenus in the chloroplast and PPC were comparable, despite a slightly lower mVenus fluorescent signal in cells expressing GES-mVenus in the PPC, indicating a possibly lower enzyme amount (Fig 3B). These observations suggest the presence of pools of GPP of comparable size in all subcellular locations, or possibly a more efficient conversion of GPP to geraniol in the PPC than in the chloroplast.

Our findings extend earlier reports that diatoms such as *P. tricornutum*^15^ and *Haslea ostrearia*^73^ maintain GPP pools in both the cytosol and chloroplast, either as intermediate compound for the biosynthesis of FPP and GGPP, or for yet unknown metabolic functions. Of the five uncharacterized, putative prenyl transferases present in *P. tricornutum*, at least one of them (Phatr3_J19000) is predicted to localize in the chloroplast, and two (Phatr3_J49325, Phatr3_47241) in the cytosol with high confidence^16,61^. The presence of GPP pools in these two locations, suggest that at least two of them have some GPP synthase activity. The remaining two (Phatr3_J15180 and Phatr3_J16615) do not have clear predictions in terms of their subcellular localization, but they are not predicted to be PPC-bound^61^. As in the case of DMAPP, the presence of a GPP pool in this compartment might suggest the presence of metabolite trafficking or of an unknown source of isoprenoid precursors.

The yields of geraniol that we recorded are the highest reported for the auxotrophic production of monoterpenoids in microalgae to date, with the cytosolic expression of GES-mVenus which was higher those previously reported for *P. tricornutum* in autotrophic growth settings (0.309 mg/L)^16^ and *C. reinhardtii* (0.100 mg/L), the latter obtained in mixotrophic conditions^62^. The differences from previously reported yields in *P. tricornutum*, could also reflect the longer cultivation time applied to this study. Our quantification might be an underestimation of the actual production capacity, as measuring geraniol and monoterpenoids production precisely in algae is a common challenge^15,62^ due to the incomplete trapping by the IM overlay, the non-optimal performance of IM as solvent for geraniol capture due the high volatility of geraniol, and its challenging separation in GC-MS/FID l^16,64^. Our results contrast with a recent report on the engineering of *P. tricornutum* with a pinene synthase (AgPinS) expressed in both cytosol and chloroplast, with products detected only the chloroplast, despite the presence of functional AgPinS in the cytosol^74^. Geraniol and other monoterpenes (e.g., limonene) can be toxic and interfere with membrane stability^76^ and have been predicted to inhibit HMG-CoA reductase (HMGR) in the MVA pathway in mammals^78,79^. If HMGR were strongly inhibited in P. tricornutum, one might expect low geraniol production in the cytosol; however, cytosolic titers were nearly twice as high as in the PPC or chloroplast (Fig. 3B), suggesting that this inhibition may be weak, non-applicable to the HMGR of *P. tricornutum*, or that geraniol rapidly escapes the cytosolic matrix.

#### FPP availability and zizaene production via heterologous ZS activity

In land plants and other photosynthetic eukaryotes with both MVA and MEP pathways, the C_15_ FPP precursor pool in the cytosol derives from the MVA pathway to supply the synthesis of essential isoprenoids such as sterols and protein farnesylation^77,78^. In these organisms an FPP pool may also be synthesized in the chloroplast by the MEP pathway to support the biosynthesis of carotenoids via its conversion to GGPP. In organisms that only possess the MEP pathway, FPP is instead produced in the chloroplast and exported to the cytosol^79,80^.

The enzyme ZS converts the FPP into the sesquiterpenoid zizaene. Diatoms transformed with ZS-mVenus constructs were cultivated with a dodecane overlay for 13 days^82^. During cultivation, all strains produced zizaene (Fig. 3C), emitted its distinctive scent. These strains also exhibited growth defects (Fig. 4C) where some cells formed aggregates, which may reflect cellular stress or compound-associated toxicity. This phenotype could arise either from accumulation of excess isoprenoid precursors, as previously reported in *E.coli* expressing heterologous terpenoid synthases^83,84^, potentially resulting from a metabolic pull caused by the increased need for FPP or, more plausibly, from the toxicity of zizaene itself. Zizaene-related toxicity has previously been reported in *E.coli* strain engineered with a polycistronic vector carrying MVA pathway genes and ZS^62,63^. This observed phenotype may reflect product toxicity, potentially due intracellular accumulation of zizaene or diffusion across the cellular membranes. To minimize these effects, we expressed the strains under the control of an inducible promoter (AP1)^55^. however, we could not detect zizaene or other related products in any of these strains (data not shown). We nevertheless decided to characterize to the best of our capacity, the strains expressing ZS-mVenus variants constitutively. Because of the formation of some cell aggregates, we could not perform flow cytometry measurements and thus determined mVenus fluorescence (Fig 3C). Instead, zizaene production was normalized to optical density (OD_750_) rather than cell number, which still provided consistent quantification across replicates, as most cells remained in suspension and a negligible amount flocculated. OD measurements were taken several times for each sample to minimize discrepancies. A prominent zizaene peak was detected all three tested compartments, verified by GC–MS in the dodecane overlay (Fig. S4), and quantified by GC–FID (Fig. 3C). Strains expressing cytosolic (0.392 ± 5.37 mg/L) and chloroplast ZS-mVenus (0.391 ± 6.10 mg/L) strains showed similar production titers, followed closely by PPC-targeted strain (0.381 ± 5.15 mg/L) (Fig. 3C). Due to the impaired phenotype, these results should be interpreted semi-quantitively. Nonetheless, they demonstrate that FPP is accessible in the cytosol, chloroplast, and PPC, and can support terpenoid biosynthesis in all three compartments. As in the case of DMAPP and GPP, also the presence of FPP in the PPC might be attributable to exchanges between cytosol and chloroplast, or less likely to uncharacterized metabolic conversions in this compartment. These results suggest that least one of the putative PTSs predicted to be resident in the cytosol (Phatr3_49325 and/or Phatr3_J47271) is capable to produce FPP, as well as at least one of the ones in the chloroplast (only Phatr3_J19000) based on computational analyses^16,61^. In *H. ostrearia*^73^ and *C. reinhardtii*^79^, FPP is found in the chloroplast as presumed earlier for the production of carotenoids. However, a recent study in *C. reinhardtii* indicates that there is no substantial reverse flux of FPP from the cytosol to the chloroplast, reinforcing the chloroplast as the promising compartment for sesquiterpenoid production in this host^85^.

**Figure 4:**
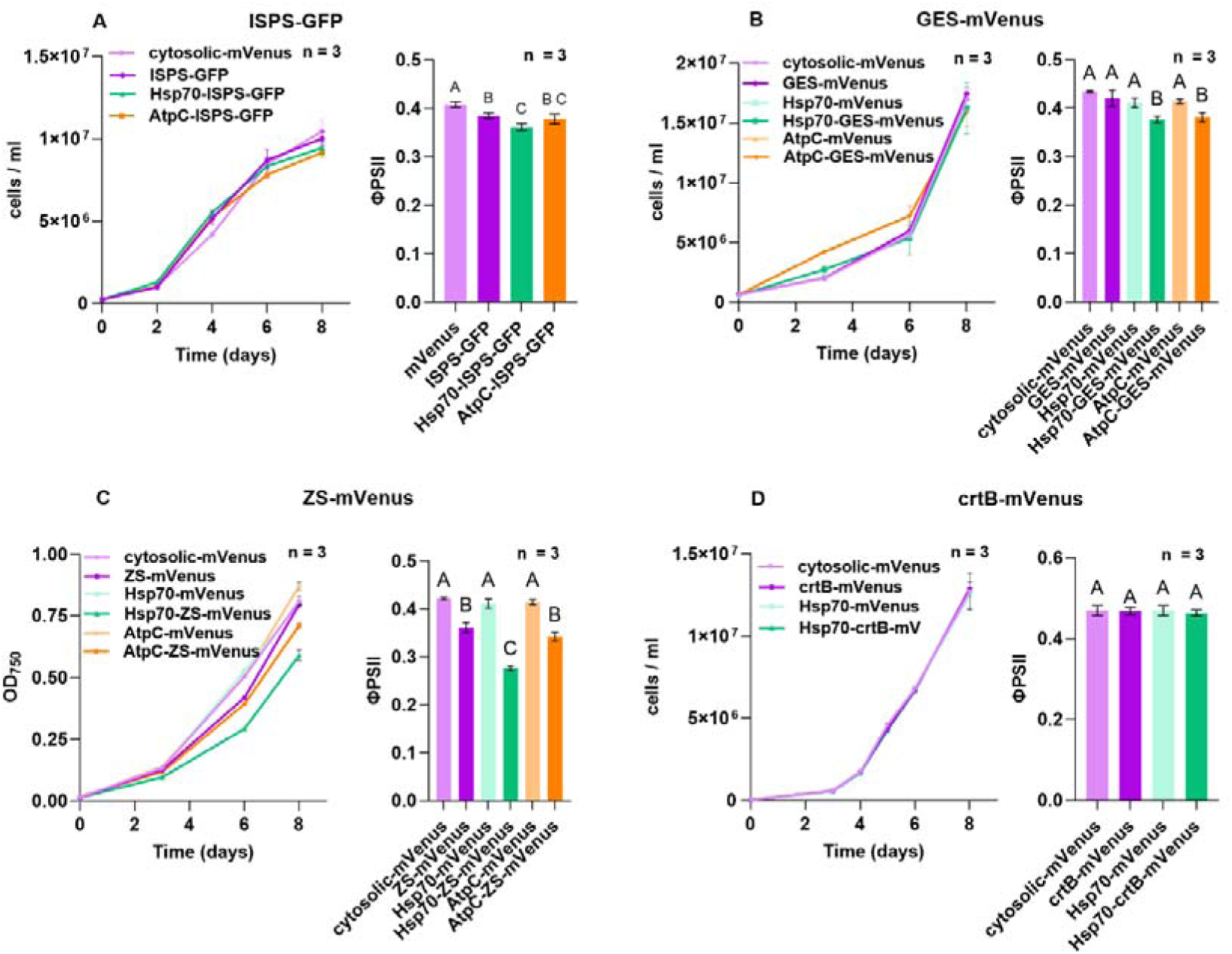
Physiological assessment of *P. tricornutum* lines expressing terpenoid synthases in different compartments. Growth data is shown on the left and effective quantum yield of photosystem 2 (ΦPSII) on the right of each panel: diatom cell lines overexpressing variants of **(A)**ISPS-GFP **(B)** GeGES-mVenus, **(C)** ZS-mVenus, **(D)** crtB-mVenus. For ZS-mVenus lines optical density (OD) was measured as an approximate estimate of growth instead of cell density via flow cytometry, as some cells produced aggregates in these cultures. Statistical differences between groups in ΦPSII graphs were calculated via one-way ANOVA (*p*-value < 0.05) and are indicated by different letters.

Our results also demonstrate that *P. tricornutum* is not an ideal host for the production of zizaene, due to induced toxicity. In contrast, other sesquiterpenes such as bisabolene have previously been heterologously produced in diatoms with no toxic effects^87^. Moreover, high volatility of zizaene has been well documented, and the use of polymeric adsorbers has been shown to improve its recovery during downstream processing^47,86^. Due to volatilization and gas exchange during cultivation, the actual yields reported in our study are likely an underestimation of the actual amounts produced. Interestingly, the only reported study on heterologous expression of the same ZS from *C. zizanioides* in *C. reinhardtii* showed no signs of toxicity, and the product was quantifiable in dodecane overlays by GC-MS/FID (26.4 fg/cell)^81.^ This highlights physiological differences between diatoms and green algae and suggests that *C. reinhardtii* may be a better production host for this compound.

#### GGPP availability and phytoene production via heterologous crtB activity

In plants and algae, the conversion of GGPP into 15-cis-phytoene is catalyzed by phytoene synthase (PSY, EC: 2.5.1.32), a plastid-localized enzyme that performs a head-to-head condensation of two GGPP molecules, releasing diphosphate^89,90^. Phytoene is normally undetectable because it is immediately converted into downstream desaturases (PDS, ZDS, EC: 1.3.99.31 and EC: 1.3.99.26, respectively) and isomerases (CRTISO, EC: 5.2.1.13), into coloured carotenoids^91^. The bacterial crtB is functional homolog of PSY that unlike plant PSY can operate ectopically when expressed in heterologous compartments such as the cytosol or non-native plastid subdomains^92,93^.

Consistent with this, overexpression of crtB-mVenus in both the cytosol and PPC resulted in detectable phytoene accumulation by HPLC (Fig. S5), with higher levels in the cytosolic variant (2.8584 ± 1.5195 µg/g DW vs 0.3793 ± 0.1608 µg/g DW) detected in the PPC (Fig. 3D). Phytoene content was normalized to dry biomass (µg/g DW), the standard metric in carotenoid quantification, as it more accurately reflects intracellular pigment composition per unit biomass with few interferences by differences in culture volume or cell density. Phytoene accumulation did not correlate with the fluorescent signal associated with the accumulation of crtB-mVenus, unlike the patterns observed for ISPS and GES. This could indicate that in this case enzyme abundance alone does not determine product yield, as metabolic context and precursor supply play a more dominant role^96^.

In land plants, cytosolic expression of crtB typically fails to yield phytoene unless co-expressed with upstream enzymes such as HMGR and crtE, which increase the MVA and GGPP supplies, respectively^97,98^. By contrast, our results in *P. tricornutum* indicate that a cytosolic GGPP supply is present and sufficient to sustain heterologous flux. In diatoms, the MVA pathway supports sterol biosynthesis and other extraplastidial isoprenoids, providing access to prenyl precursors outside the plastid^19,20,29,73^. In some species such as the diatom *H. ostrearia*, cytosolic GGPP can also be formed locally by a cytosolic GGPPS isoform^73,99^. In addition to local synthesis, the cytosol may also receive IPP, DMAPP or short-chain prenyl diphosphates exchanged between compartments, particularly when demand increases^73,99^. The observed differences in phytoene accumulation between compartments likely reflect both substrate availability and the capacity to accommodate hydrophobic intermediates. In the plastid of plants and algae, PSY has access to abundant MEP-derived GGPP however, phytoene is rapidly consumed by downstream enzymes, which generally prevents its accumulation to detectable levels in green tissues^94,100^. As for other prenyl phosphates detected in the PPC, GGPP likely derives from the exchange with the plastid or cytosol, or, less likely from unknown enzymatic source, as no conventional prenyl transferase-based pathway operates in this compartment. This restricted precursor supply likely limits the amount of phytoene that can be produced. In the cytosol, however, expression of crtB can draw on GGPP supplied by the MVA pathway and/or precursor exchange without the tight regulatory constraints that limit plastidial PSY activity, which in plants include feedback inhibition by downstream carotenoids and post-translational control through interactions with regulatory proteins such as ORANGE and Clp protease systems^101^ resulting in more robust phytoene accumulation^89,93^. Differences may also be related to the physical capacity of each compartment: cytosolic production benefits from extraplastidial sinks such as endomembrane and lipid droplets that can sequester carotenoid intermediates without impairing photosynthesis^97,102,103^, whereas the PPC may provide only limited storage space, which could restrict the extent of phytoene accumulation despite the presence of substrate.

### Effects of compartmentalized heterologous terpenoid biosynthesis on growth and photosynthetic performances

Occasionally, the accumulation of bioactive terpenoids is known to cause cellular toxicity, often due to their cytotoxic, membrane-disrupting, or antimicrobial properties^109^. Additionally, heterologous terpenoid biosynthesis relies on the endogenous pools of IPP and DMAPP, and perturbations in their biosynthesis and availability may disrupt the homeostasis of essential metabolites such as sterols, pigments, and other compounds involved in cell physiology. We hypothesized that compartmentalizing terpenoid biosynthesis into organelles such as the PPC could mitigate toxicity or reduce metabolic competition. Although isoprene, geraniol and zizaene are volatile and can escape the cellular membranes, we set out to evaluate whether the heterologous terpenoid production in different subcellular compartments differentially affected diatom physiology, by profiling growth and photosystem II quantum yield (ΦPSII) during 8 days of batch cultivation in cell lines expressing ISPS-GFP and CrtI-mVenus constructs, and 13 days for cell lines expressing GES-mVenus and ZS-mVenus.

Most transformants did not show altered growth performance. All ISPS-GFP strains were comparable to the cytosolic mVenus control (Fig. 4A). Although isoprene emission has been linked to improved thermal tolerance and photosynthetic stability in higher plants^75,76^, our data shows a modest reduction in ΦPSII in *P. tricornutum* expressing ISPS-GFP (Fig. 4A). This species naturally produces only trace amounts of isoprene^77^. The observed reduction of ΦPSII in Hsp70-ISPS-GFP is minor and of uncertain physiological relevance. GES-mVenus transformants also showed no significant difference in growth compared to the mVenus control^16,64^. As reported here, the accumulation of geraniol does not affect the algal growth in line with the previous reports shown in *P. tricornutum*^16^ and *C. reinhardtii*^64^. The ΦPSII was same as the control in cytosolic lines in line with previous reports in *C. reinhardtii*^64^ but strains expressing GES-mVenus in the chloroplast and PPC display lower maximum quantum yields of photosystem II (Fig. 4B). For AtpC-GES-mVenus expressing cell lines, this reduction may reflect competition for GPP pools in the chloroplast which may potentially interfere with carotenoid biosynthesis hence, yielding lower ΦPSII. A decreased carotenoid supply can compromise photoprotection and light-harvesting efficiency, leading to reduced photosynthetic performance under high-light conditions^110^. A similar explanation may apply to strains expressing GES-mVenus in the PPC, particularly if the GPP converted to geraniol is trafficked towards the chloroplast to supplement its internal pool of prenyl phosphates for carotenoid biosynthesis. Alternatively, geraniol produced in the chloroplast or in the PPC, which surrounds the chloroplast, may accumulate at higher local concentrations in proximity to the thylakoid membranes, potentially affecting their stability more than geraniol synthesized in the cytosol. Overall, geraniol production did not cause major growth defects on the *P. tricornutum* cells.

All zizaene producing strains displayed transition to the oval morphotype and pronounced cell aggregation suggesting product toxicity or a metabolic burden specific to this sesquiterpene. The localization of ZS-mVenus in the PPC had most pronounced growth inhibitory effects, followed by strains producing zizaene in the chloroplast and cytosol (Fig. 4C). Across all compartments, zizaene-producing strains showed reduced photosynthetic efficiency (ΦPSII), with the lowest quantum yield in photosystem II in PPC-targeted strains (Fig. 4C)^59,60^. These results, although more pronounced, mirrored the impact observed for geraniol production in the PPC and chloroplast on ΦPSII.

Diatom cell lines expressing crtB in either the cytosol or PPC showed no differences in growth relative to mVenus or HSP-mVenus controls. This suggests that phytoene production does not impose a detectable metabolic burden even if this may be due to low amounts of heterologous product. Importantly, phytoene accumulation had no measurable impact on photosynthetic activity in any compartment (Fig. 4D). Unlike isoprene, geraniol, and zizaene, phytoene is non-volatile and therefore remains confined to the compartment in which it is synthesized. Moreover, phytoene is endogenously produced in the chloroplast of *P. tricornutum*, making it less likely to exert toxic effects. This mirrors findings in plants, where cytosolic phytoene accumulation minimally affects photosynthesis^97,103^. Even when phytoene accumulated in the PPC, ΦPSII remained unchanged. This observation supports the idea that the PPC functions as a cytosol-like compartment, largely insulated from the chloroplast. However, it is also possible that phytoene levels in the PPC were simply too low to exert any disruptive effect on photosynthesis, especially considering that measurable impacts on photosynthetic activity in plant leaves have only been reported at phytoene concentrations far higher than those attainable in the PPC^97^.

These findings indicate that PPC compartmentalization does not inherently prevent toxicity when compounds are volatile and capable of crossing membranes. In contrast, non-volatile terpenoids such as phytoene appear well tolerated, suggesting that confinement within the PPC can be an effective strategy only for metabolites that remain localized.

### Metabolic and physiological implications of the presence of prenyl phosphates in all three subcellular compartments

Our results demonstrate the presence of DMAPP, GPP, FPP, and GGPP in the cytosol, chloroplast and surprisingly in the PPC of *P. tricornutum*. As no prenyl transferases are predicted to localize to the PPC, we hypothesize that prenyl phosphates accumulate in the PPC as a result of metabolic trafficking rather than unconventional PPC-bound biosynthesis. Some studies suggest that *P. tricornutum* does not exhibit full prenyl phosphate crosstalk between cytosol and chloroplast^75^; however, the detection of these prenyl phosphates in the PPC supports at least partial metabolite exchange, which could be driven by either active or passive transport. Dedicated isoprenoid transporters may exist, and prenyl phosphates could also be exchanged across organelles via membrane contact sites (MCS’) or, less likely, by direct diffusion trough the membranes. In these scenarios, *P. tricornutum* may transfer prenyl phosphates constitutively, or only under specific conditions. Another possibility could be that the metabolic exchange is restricted to cytosol–PPC or chloroplast– PPC connections. It is also possible that the red algal ancestor involved in the secondary endosymbiotic event giving rise to diatoms already possessed mechanisms for prenyl phosphate exchange, which were retained to facilitate metabolite flow between compartments^112^.

The presence of GGPP pools in the cytosol of *P. tricornutum* is particularly intriguing, as no tetraterpenoids or other higher isoprenoids are currently known to be synthesized in this compartment in *P. tricornutum*. One explanation is that cytosolic prenyl transferases, which often exhibit relaxed substrate specificity, may generate GGPP as a side product while elongating shorter prenyl phosphates. Among the five putative prenyl transferases annotated in *P. tricornutum*, two are predicted to localize to the cytosol and could therefore contribute to GGPP formation. Another possibility is metabolic crosstalk with other organelles in the cells, whereby GGPP or its precursors are exchanged from organelles where this metabolite is required, such as mitochondria, where GGPP serves as a precursor for ubiquinone biosynthesis^113^. Structural studies in other organisms have shown that prenyl diphosphate synthases can produce products of variable chain length depending on active-site architecture and substrate availability^77–79^. Thus, the presence of GGPP in the cytosol may not necessarily reflect a dedicated biosynthetic pathway for tetraterpenoids, but rather the combined outcome of side-product formation, inter-organelle exchange, and metabolic flexibility. Importantly, even if not used for carotenoid biosynthesis, cytosolic GGPP could fulfill essential roles in post-translational protein modification such as geranylgeranylation, thereby linking its presence to core cellular functions.

In this context, the PPC can provide a unique experimental window into inter-pathway communication that remains poorly accessible in plants. In fact, diatoms, with their compartmentalized metabolism and genetic tractability, represent an exceptional model to investigate this crosstalk and address long-standing questions about isoprenoid precursor transport and coordination. Another factor under consideration is the mildly acidic pH of PPC (∼6.8)^66^, which contrasts with the more alkaline cytosol (∼7.9) and stroma (∼8.0), which is maintained and tightly linked to the diatom carbon-concentrating mechanisms^65^. The acidic pH imposes constraints on pathway design, but modest, localized adjustments together with PPC-targeted enzymes that function well around pH 6.8-7 remain feasible strategies for PPC engineering. Furthermore, our results highlight that the prenyl phosphate precursors in the PPC can be utilized for the synthesis of heterologous terpenoids. From a metabolic engineering perspective, although the PPC may not be optimal for the synthesis of volatile terpenoids such as isoprene, geraniol and zizaene, it offers clear potential for non-volatile products like phytoene which requires a physical isolated storage space where they can be sequestered from the central metabolism. Compounds such as phytoene or squalene, normally directly channelled in their respective downstream pathways (i.e., carotenoids and sterols, respectively), could find in the PPC a suitable accumulation space.

## Conclusions

In this study, we provided the proof-of-concept of the full spectrum potential of *P. tricornutum* for the biosynthesis of heterologous terpenoids in the cytosol, chloroplast and PPC, due to the availability of all main prenyl phosphates in all three compartments, highlighting the high metabolic versatility of this organism for heterologous terpenoid production. Importantly, we demonstrated that the PPC can be repurposed as a functional, dedicated subcellular hub for heterologous terpenoid biosynthesis isolated from the resto of the cell. DMAPP, GPP, FPP and GGPP pools are available in the PPC of *P. tricornutum* in amounts sufficient to support the heterologous biosynthesis of hemi-, mono-, sesqui- and tetraterpenoids. As there is no endogenous biosynthetic pathway predicted in the PPC^39^, these results provide strong indirect evidence for precursor trafficking between the MVA and MEP by active transport or passive diffusion into/through the PPC or less likely, of unknown biosynthetic processes. The crosstalk between MEP and MVA pathways has long been proposed in plants but remains poorly understood, and the diatom PPC of diatoms acting as an interface will serve as an additional valuable tool to investigate this mechanism. Our work provides the first systematic profiling of compartment-specific terpenoid biosynthetic capacity of a diatom revealing distinct capacity of the production of terpenoids deriving from the main prenyl phosphate precursors across compartments. Monoterpenoid production (geraniol) was most efficient in the cytosol, yielding up to 1.85 mg L^−1^, representing the highest geraniol titer reported in microalgae to date, sesquiterpenoid production (zizaene) was at the same level in all compartments, while hemiterpenoid production (isoprene) was favored in the chloroplast and tetraterpenoid (phytoene) was efficiently produced in cytosol. In this study, in non-optimized conditions, the PPC yielded the lowest titers for all the terpenoids, compared to the cytosol or chloroplast. These patterns likely reflect differences in prenyl phosphate precursor pools, as well as compartment-specific enzyme activity and concentrations. Importantly, tagging heterologous terpene synthases to the PPC did not generally induce major growth defects even though lower ΦPSII could be observed with different severity in cell lines expressing ISPS-GFP, GES-mVenus, and ZS-mVenus targeted to PPC, indicating that the PPC can accommodate heterologous metabolic flux.

Beyond its current functionality, and envisioning pathway optimization strategies that include the maximization of precursors availability, we anticipate that the diatom PPC represents a promising, engineerable space for synthetic biology applications. It could serve as a dedicated production organelle for modular assembly of multienzyme complex pathways. The periplastidial membrane (PPM) could also be exploited for functional expression of membrane-bound cytochrome P450s and other scaffold decorating enzymes. Further studies are required to understand the mechanism by which prenyl phosphate precursors are made available in the PPC by identifying and characterizing relevant transporters, whose manipulation might enable increased metabolic flux into the PPC and boost product titers. Taken together, our findings highlight the PPC as a promising and underexplored compartment for terpenoid production in diatoms.

Finally, the ability to target and assemble complex metabolic pathways in this compartment, makes it a versatile minimal subcellular space directly coupled with a eukaryotic photosynthetic chloroplast, which could be suitable for specialized functions, opening new synthetic biology opportunities the optimized production of heterologous terpenoids or other applications in diatoms.()Supporting information()Figures S1–S5 contain additional experimental data including control microscopy images, chromatograms and mass spectra, and Table S1-S3 with details on constructs, primers sequence and primers used in this study.

## Author Contribution

MF conceived the study, with contributions from PP, FP, LM; PP, FP, LM and MF designed the experiments; PP, FP and LM carried out the experiments. PP, FP, LM, MF analyzed and interpreted the data. PP wrote the manuscript with contributions from FP, LM and MF. All authors read and approved the manuscript.

## Acknowledgments

This work was supported by VILLUM FONDEN grant to MF (grant number 3721), by a SDU Climate Cluster (SCC) Research Infrastructure Grant to MF. LM was supported by HORIZON-MSCA-2023-PF-01, Project: 101149868. We thank Maxime Bastide for his contribution to geraniol extractions. We are thankful to Lars Duelund for the help in GC-MS measurements.

## List of abbreviations

AP1: alkaline phosphatase
AtpC: γ subunit of the thylakoid ATP synthase
crtB: phytoene synthase
DMAPP: dimethylallyl pyrophosphate
GFP: green fluorescent protein
GGPP: geranylgeranyl diphosphate
GES: geraniol synthase
GPP: geranyl pyrophosphate
Hsp70: heat shock protein 70
IDI: isopentenyl diphosphate isomerase
ISPS: isoprene synthase
MVA: mevalonate pathway
MEP: methylerythritol 4-phosphate pathway
PPC: periplastidial compartment
PPM: periplastidial membrane
PSY: phytoene synthase
PDS: phytoene desaturase
wt: wild type
ZS: zizaene synthase.

