## Supplementary information file 1 for "Repurposing the diatom periplastidial compartment for heterologous terpenoid production"

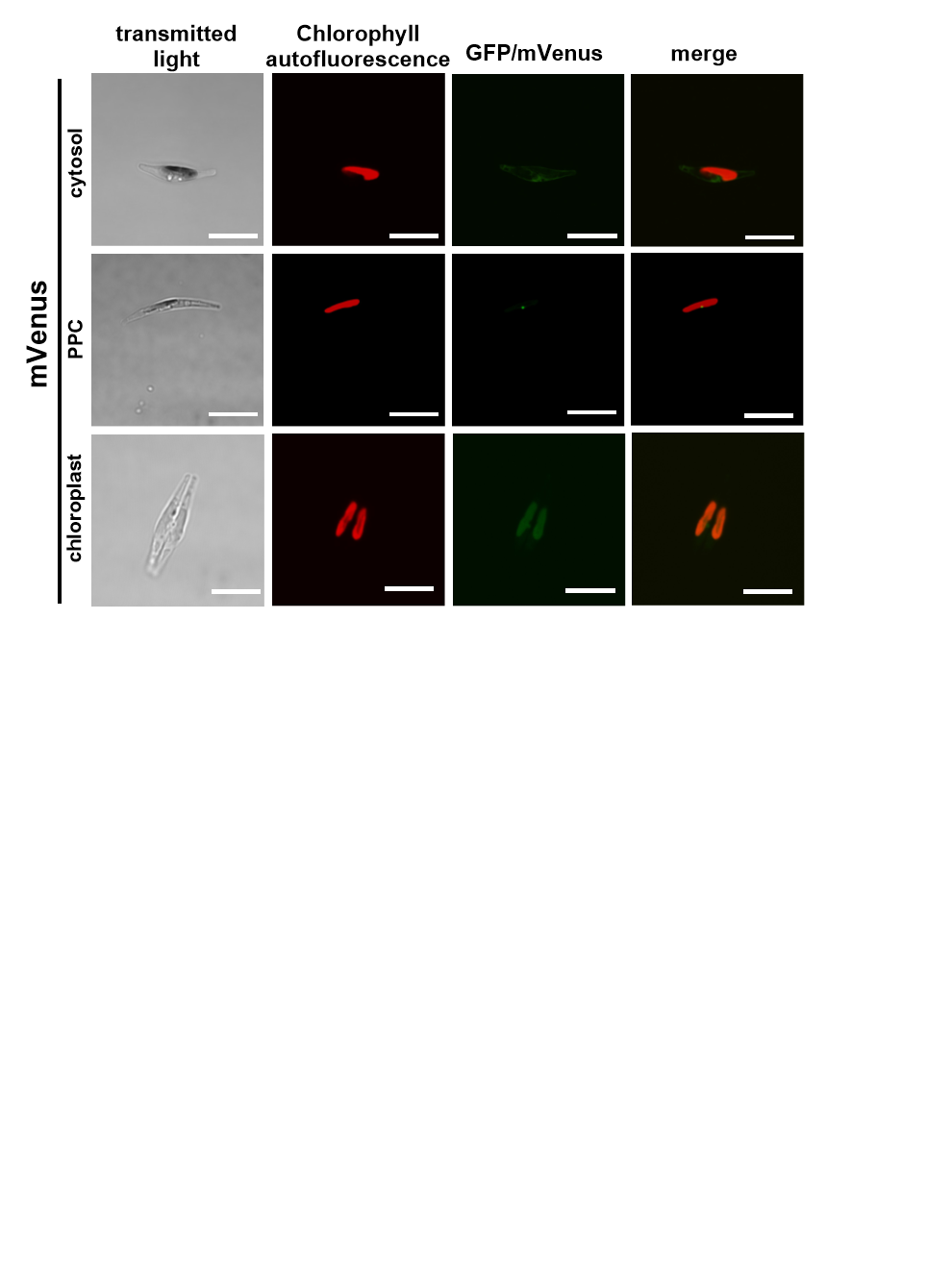

**Figure S1**. **Subcellular localization of recombinant mVenus targeted to the cytosol, PPC and chloroplast in *P. tricornutum*, as control cell lines for the subcellular localization of heterologous terpene synthase (Fig.1).** Each image is representative of at least three independent cell lines per construct. Scale bars represent 5 µm

.

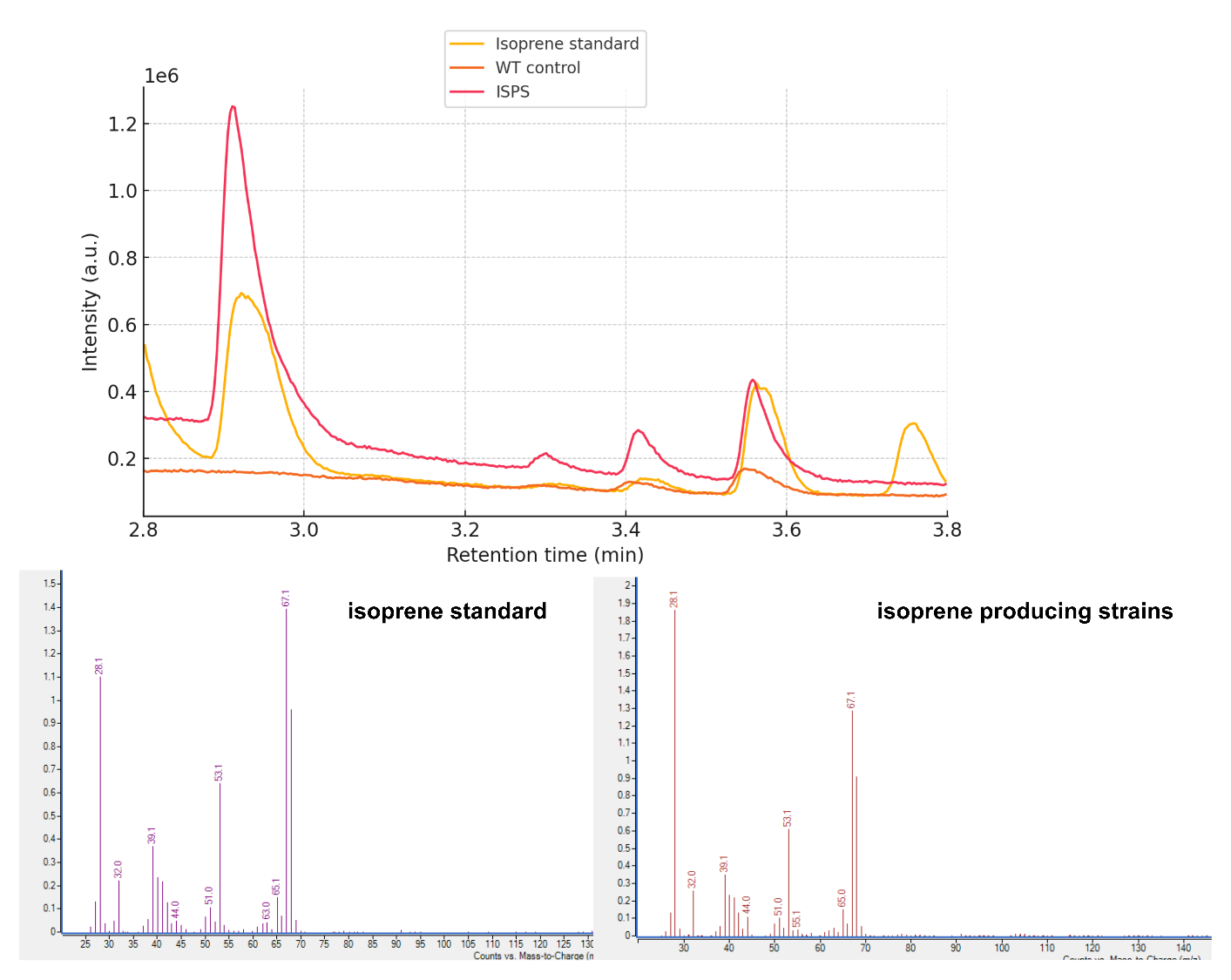

**Figure S2:** Representative chromatograms of isoprene (RT: 2.9 min) and mass spectra, from transgenic diatom and wild-type strains, compared to the authentic standard.

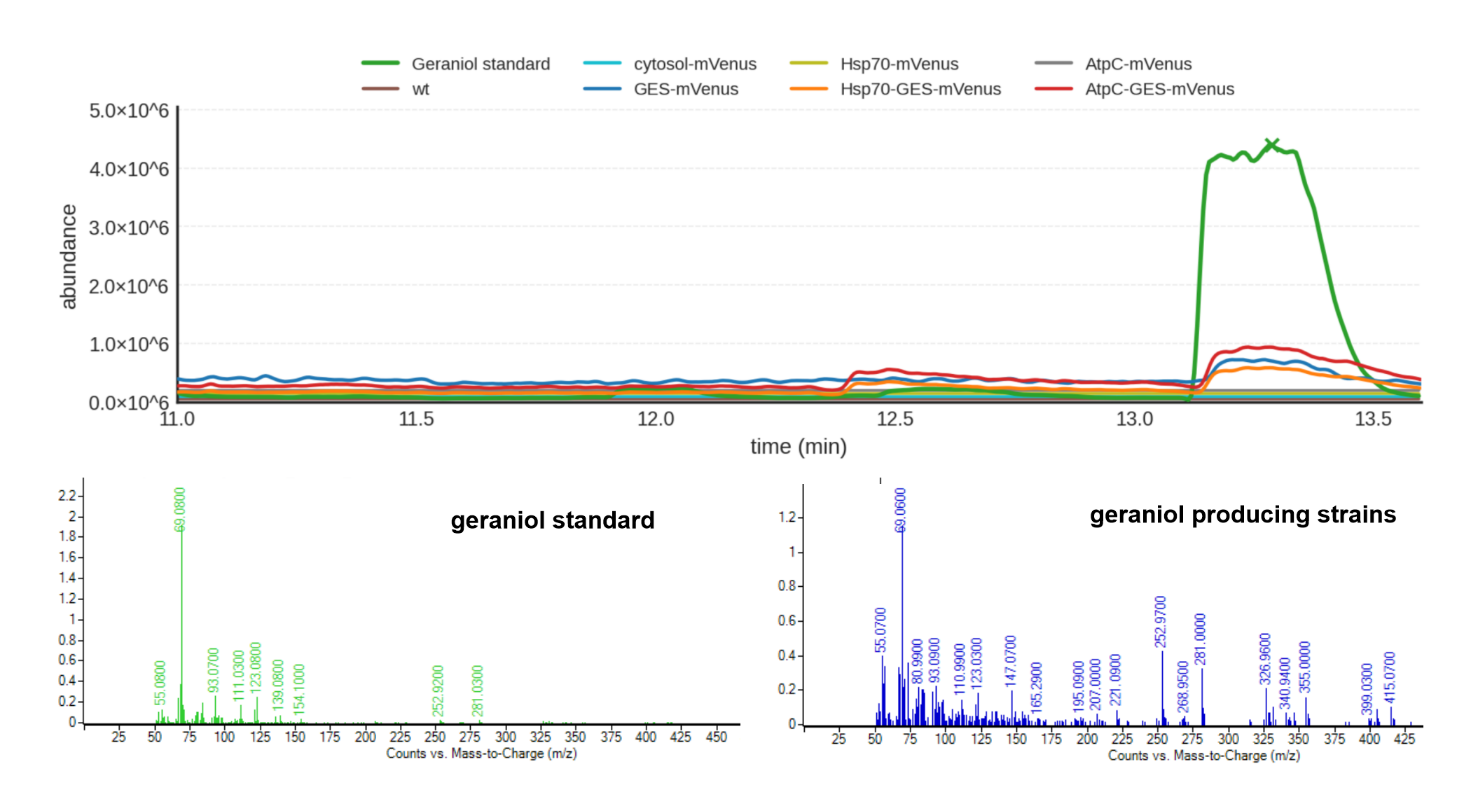

**Figure S3:** Representative chromatograms of geraniol (RT: 12.3 min) and mass spectrum, from transgenic diatom and wild-type strains, compared to the authentic standard.

Supplementary Figure 4: Representative chromatograms and fragmentation of zizaene producing strains and cedrene standard.

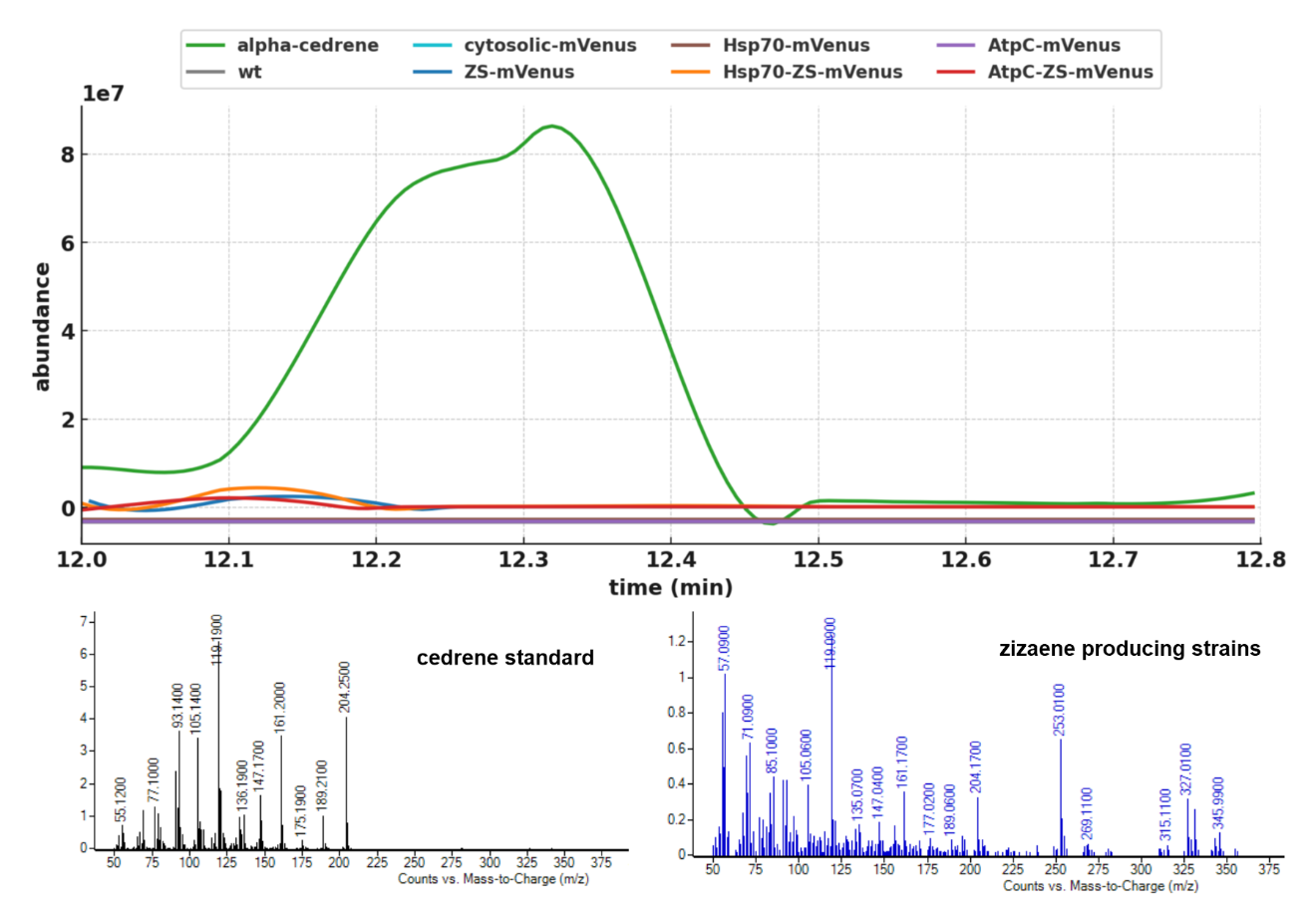

**Figure S4:** Representative chromatograms of zizaene (RT: 12.3 min) and mass spectra, from transgenic diatom and wild-type strains, compared to the cedrene authentic standard. Cedrene was employed as standard due to its close similarity to zizaene, whose commercial standard is not available.

.

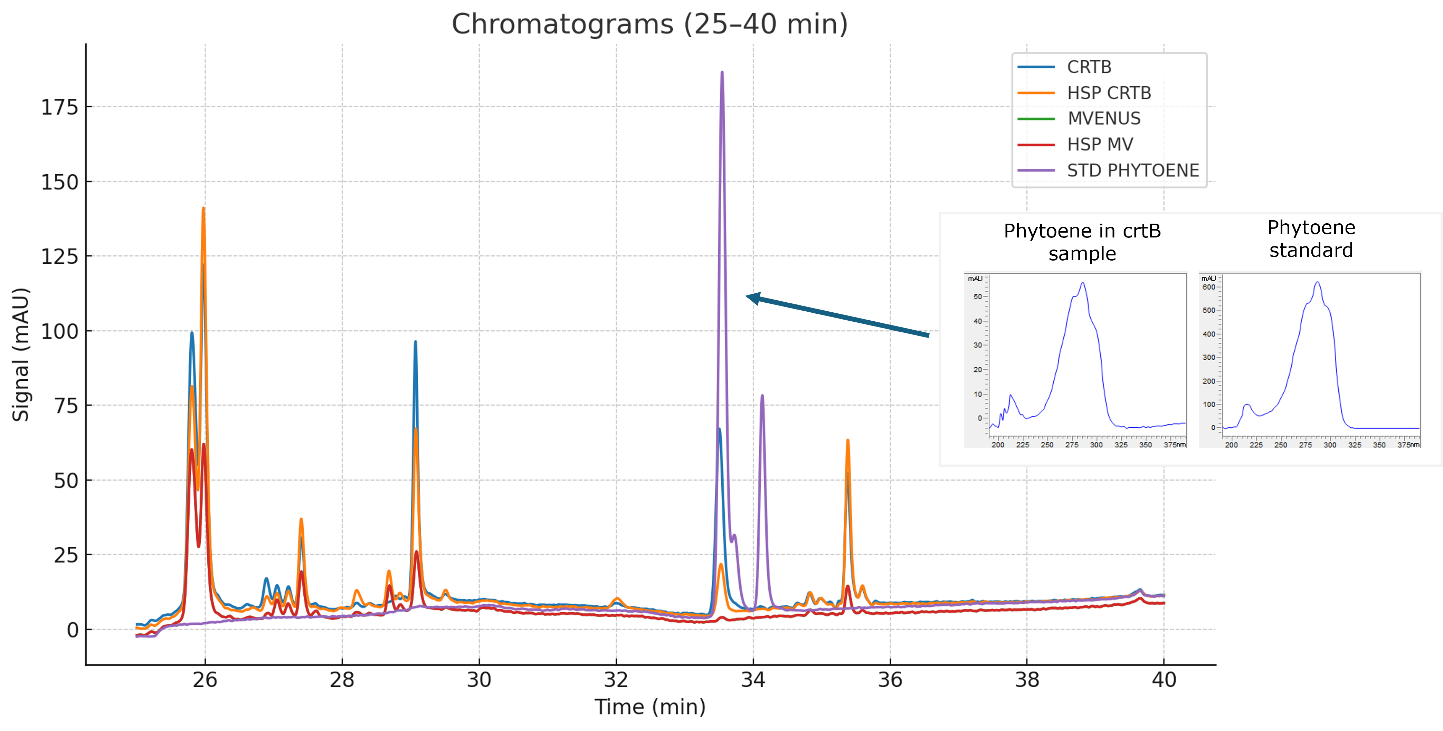

**Figure S5.** Representative chromatograms and fragmentation of phytoene producing strains and UV spectra of heterologous phytoene and phytoene authentic standard.

**Table S1**: uLoop used for the assembly of constructs

| **Syntax /Part** | **Gene ID** | **Description** | **Reference** |
| --- | --- | --- | --- |
| L1-1 PTCv2 | N/A | Conjugation and prop. Elements | (Pollak et al., 2019) |
| L0_AC_pPt49202 | *Phatr3_J49202* | Promoter | (Pollak et al., 2019) |
| L0_AC_pPtAP1 | *Phatr3_J49678* | Alkaline phosphatase promoter | (Lin et al., 2017) |
| L0_EF_tPtJ25172 | *Phatr3_J25172* | Terminator (*FcpB*) | (Pollak et al., 2019) |
| L0_DE_mVenus | mVenus-stop | Fusion tag | (Pollak et al., 2019) |
| L0_DE_3xstopcodon | - | stop codon | (Pollak et al., 2019) |
| L0_HSP70_transit peptide | *Phatr3_J21519* | Hsp70 transit peptide | (Apt et al., 2002) |
| L0_AtpC_transit peptide | *Phatr3_J20657* | AtpC transit peptide | (Kilian et al., 2005) |
| L0_CD/DE_ISPS | ABV04402.1 | Isoprene synthase | This study |
| L0_CD/DE_GES | JN882024.1 | Geraniol synthase | This study |
| L0_CD/DE_ZS | HI931360.1 | Zizaene synthase | This study |
| L0_CD/DE_crtB | ADD79329 | Phytoene synthase | This study |

**Table S2**. Primers used

| **Primer ID** | **Sequence** |
| --- | --- |
| UNS1F | CATTACTCGCATCCATTCTC |
| UNSXR | CTTGGGAAGATCGTAGTGTA |

**Table S3.** Coding sequences (included between the 5’ and 3’ of the uLoop overhangs) of the L0 parts generated in this study.

**L0_Hsp70_N-terminal_BTS 165 bp**

GTCCACCTTCCTTCCAGTTCGACTCTGTTGACCTGCGTGTCCGTCCTCTTGTCCGGAGCCCACCCGGCGAAAGCCTCTTGGCTCGCCCGTCGAACGGTTGAAAAGCCCACCCTCGCTCGTATTCATGAGCAGCGCGACAGCACAGATCGCAAGTCGCGGGCACCC

**L0_AtpC_N-terminal_BTS 132 bp**

CGTTCGTTCTGCATTGCCGCGCTCTTGGCAGTTGCCTCCGCTTTTACTACGCAGCCGACATCCTTCACCGTGAAGACCGCTAACGTCGGAGAACGCGCCAGCGGTGTCTTTCCCGAGCAATCTAGTGCCCAC

**L0_ISPS-GFP_DE bp**

GAAAAGGCCGAATTCTTGACCCTCCTCGAACTTATTGATAACGTGCAGCGCCTTGGGTTAGGATACCGTTTCGAGTCCGACATCCGGGGAGCTCTTGACCGCTTCGTCTCCTCTGGAGGCTTCGACGCAGTCACCAAGACAAGTTTGCACGGGACGGCCCTAAGTTTCCGTCTGTTGCGCCAACATGGATTTGAGGTCAGCCAGGAAGCCTTTTCCGGATTTAAGGACCAGAATGGTAATTTCCTCGAAAATTTGAAGGAAGACATTAAGGCGATTCTTTCCCTTTACGAAGCCTCATTCCTCGCCCTCGAAGGTGAAAATATCCTCGATGAAGCTAAGGTATTCGCCATCTCGCATCTGAAGGAATTGAGCGAAGAGAAGATTGGCAAGGAACTTGCTGAACAAGTCAATCATGCCCTAGAGCTCCCCCTGCACCGTCGTACCCAACGACTCGAAGCAGTGTGGTCGATCGAAGCATACCGTAAGAAAGAAGACGCCAACCAGGTTCTTTTGGAACTGGCCATCTTGGACTACAACATGATTCAATCCGTCTACCAGCGTGATCTCCGTGAAACCTCCCGCTGGTGGCGCCGCGTCGGACTCGCTACCAAGCTCCACTTTGCCCGCGATCGGTTGATTGAGTCTTTCTACTGGGCGGTGGGCGTAGCGTTCGAGCCCCAGTACTCCGACTGTCGTAACTCAGTCGCTAAGATGTTTTCCTTCGTCACTATTATTGACGATATCTACGACGTCTATGGTACCCTGGACGAGCTCGAACTTTTCACGGATGCGGTCGAACGTTGGGATGTCAACGCAATTAACGACCTCCCCGACTACATGAAATTGTGCTTTCTCGCCTTGTACAACACGATCAACGAGATAGCGTACGATAACCTCAAGGACAAGGGCGAGAACATCCTGCCGTACCTGACCAAAGCCTGGGCGGATCTCTGCAACGCCTTTTTGCAAGAAGCTAAGTGGCTCTATAATAAGTCGACCCCTACTTTTGACGACTACTTTGGTAACGCATGGAAGTCTTCAAGCGGTCCCCTCCAGTTGGTCTTTGCCTACTTTGCCGTCGTTCAGAACATTAAGAAAGAAGAAATTGAAAACCTTCAAAAGTACCACGATACGATCAGCCGACCCTCCCACATCTTTCGCTTGTGCAATGATTTGGCCTCTGCCTCCGCTGAGATTGCCCGCGGTGAAACCGCGAACTCGGTGTCGTGCTATATGCGTACTAAAGGTATTTCTGAGGAACTTGCCACGGAATCGGTGATGAACCTCATCGATGAAACTTGGAAGAAGATGAACAAGGAGAAACTTGGAGGATCCCTCTTCGCCAAGCCGTTCGTTGAGACAGCCATCAACTTGGCCCGTCAGAGCCACTGCACTTACCACAACGGCGACGCCCATACGAGTCCTGACGAGTTGACTCGAAAGCGAGTGCTTTCCGTCATCACAGAGCCAATTCTCCCGTTCGAACGTTCGAAGGGCGAAGAATTGTTTACGGGTGTAGTACCCATTCTTGTTGAACTCGACGGAGATGTCAACGGCCACAAGTTCTCCGTCTCTGGAGAGGGTGAAGGAGACGCCACTTACGGGAAGTTGACGCTTAAGTTTATCTGTACAACTGGAAAGCTCCCAGTTCCTTGGCCGACTCTTGTTACGACCTTGACCTACGGTGTCCAGTGCTTTGCTCGCTATCCGGATCATATGAAGCAACATGACTTCTTCAAGTCCGCTATGCCCGAAGGCTACGTGCAAGAACGGACCATTTTCTTCAAGGACGATGGCAACTACAAGACCCGAGCCGAAGTCAAATTCGAAGGGGACACCCTCGTCAATCGTATCGAACTGAAAGGCATCGACTTCAAAGAGGACGGCAATATTTTGGGTCACAAGCTCGAATACAACTACAATAGCCACAAGGTGTACATTACCGCCGATAAACAGAAGAACGGTATCAAGGTGAACTTTAAAACCCGTCACAACATCGAGGACGGTAGTGTCCAGCTCGCCGACCATTACCAACAGAACACTCCCATTGGAGATGGACCCGTCCTCCTCCCTGATAACCACTACCTGTCCACACAGTCAGCGTTGTCGAAGGATCCGAATGAGAAGCGCGACCACATGGTTCTTCTAGAGTTTGTCACCGCCGCAGGAATTACGTTGGGTATGGACGAACTGTATAAGGGGAGCGGGAGCGGGAGCGGGAGCAGGAGC

**L0_GES-mVenus_DE 2481 bp**

GCCGCTACGATCAGTAACTTGTCGTTTTTGGCTAAGTCACGCGCATTGTCTCGCCCTTCTTCTAGTTCGCTTTCTTGGCTTGAACGCCCGAAAACTTCGTCAACTATCTGTATGTCCATGCCCTCAAGTTCCTCGTCTAGTTCATCTTCCTCTATGTCCCTTCCACTTGCAACACCTCTCATTAAAGACAACGAATCACTTATTAAGTTTCTCAGACAACCCTTGGTTCTCCCTCATGAGGTAGACGACTCGACAAAACGTCGAGAGTTGTTGGAGAGAACCCGCAAAGAATTGGAGCTTAACGCCGAAAAACCGTTGGAGGCACTCAAAATGATCGACATCATCCAGAGACTCGGACTTTCGTATCACTTTGAGGATGATATCAATTCAATCCTTACAGGCTTTAGTAACATTTCGAGTCAGACGCACGAAGACCTCTTGACTGCTAGCCTCTGCTTCAGACTCTTGCGACACAACGGCCATAAGATTAATCCAGATATTTTCCAAAAATTTATGGACAATAACGGAAAGTTCAAAGATTCGCTTAAGGACGACACATTGGGTATGCTCTCGCTCTACGAAGCTTCGTATCTCGGAGCAAACGGCGAGGAGATCCTTATGGAGGCTCAAGAGTTTACAAAGACCCACCTTAAAAATTCGTTGCCAGCAATGGCTCCATCTCTTTCAAAAAAGGTTAGCCAAGCACTTGAACAGCCCCGCCACCGTCGTATGCTCCGCCTCGAGGCAAGGCGATTTATCGAAGAATATGGGGCGGAGAACGACCATAATCCCGACTTGCTTGAATTGGCCAAGCTTGATTACAATAAGGTTCAAAGCTTGCACCAGATGGAGCTTAGTGAGATCACTCGCTGGTGGAAACAGCTCGGGTTGGTCGACAAACTTACGTTCGCCAGGGATCGACCGCTTGAATGCTTTCTCTGGACCGTAGGGTTGCTTCCAGAGCCCAAGTATAGTGGTTGCAGAATCGAACTTGCAAAAACCATTGCCATTCTTCTTGTCATTGACGATATTTTCGATACGCACGGCACCCTTGATGAATTGTTGCTCTTTACGAATGCGATCAAGAGATGGGACCTTGAAGCGATGGAAGATCTCCCAGAGTATATGAGGATCTGCTATATGGCCCTTTACAATACGACAAACGAAATTTGCTATAAAGTATTGAAGGAAAACGGTTGGTCGGTCCTTCCGTATCTTAAAGCGACGTGGATTGACATGATCGAGGGGTTTATGGTTGAAGCTGAATGGTTTAACTCCGATTACGTCCCCAATATGGAAGAATACGTGGAGAACGGAGTGCGAACCGCAGGATCTTACATGGCGCTTGTCCACTTGTTTTTCCTCATTGGCCAAGGCGTCACTGAGGATAACGTGAAACTTCTCATTAAGCCCTATCCAAAGCTTTTTAGCAGCTCAGGACGTATTTTGCGACTCTGGGATGATTTGGGTACGGCTAAGGAAGAACAGGAACGCGGAGATCTCGCTTCCTCAATCCAACTTTTTATGAGGGAGAAAGAGATTAAGTCAGAAGAGGAGGGCAGAAAGGGGATCTTGGAGATCATTGAGAACCTCTGGAAAGAGCTTAACGGTGAACTTGTATACCGCGAGGAGATGCCGCTTGCGATCATCAAAACGGCATTCAATATGGCCCGTGCTTCACAGGTGGTTTATCAACACGAAGAGGACACTTATTTTAGTTCTGTTGACAACTATGTAAAAGCATTGTTTTTCACGCCCTGCTTCATGGTCAGCAAGGGTGAGGAACTCTTTACTGGAGTCGTGCCAATCCTTGTTGAGCTTGACGGAGACGTGAACGGTCACAAGTTTTCCGTTAGCGGCGAAGGAGAAGGCGACGCCACATACGGAAAGTTGACTCTCAAGCTCATCTGCACGACGGGAAAGTTGCCGGTCCCGTGGCCCACCCTCGTCACCACCCTAGGCTACGGTTTGCAGTGTTTCGCCCGCTACCCTGACCACATGAAGCAGCATGATTTCTTTAAATCGGCCATGCCCGAAGGTTATGTCCAGGAGCGTACCATTTTCTTCAAAGACGATGGTAACTACAAGACCCGCGCTGAAGTCAAGTTTGAGGGCGACACGCTGGTAAATCGAATCGAATTGAAGGGAATCGATTTCAAGGAAGACGGCAACATTTTGGGACACAAGTTGGAATACAACTATAACTCCCACAACGTGTACATCACCGCTGACAAGCAGAAGAACGGCATCAAGGCTAATTTTAAGATCCGTCACAATATTGAAGACGGCGGTGTACAGTTGGCAGACCACTACCAACAGAACACGCCCATTGGCGACGGACCGGTCCTCTTACCGGACAACCACTATCTGTCCTACCAGAGTGCCCTCTCCAAAGACCCCAATGAGAAACGTGATCACATGGTGCTGTTGGAATTCGTTACCGCTGCAGGGATCACTCTAGGTATGGATGAACTCTATAAG

**L0_ZS-mVenus_DE 2382 bp**

ATGGCGACGACTGCCGCCTTCTGCCTCACCACCACTCCGATCGGCGAACCCGTCTGCCGCCGTCAGTACCTCCCAACCGTCTGGGGCTCCTTCTTCCTCACCTACCAGCCCTGCACGCCGGAAGAGGTCCAGTCCATGGAAGAGAGGGCCCTGGCCAAGAAGACGGAGGTGGGCCGCATGTTGCAGGAGGTCGCCGCCTCCAGTAACCTCGCCCGGAAGCTGGGCCTTGTCGATGAACTCGAACGCCTCGGTGTGGACTATCACTACAAGACGGAAATCAACGACTTGCTTGGTGCCATTTACAATGGCAAGGACGACGATAACGGAGGTTCTGATGACGACCTCTACATCACATCGCTTAAGTTCTACCTCCTCCGAAAGCACGGATACGCTTTGTCTTCCGACGTCTTTCTCAAGTTCCGCGATGAGCAAGGAAATATTTCGTCGGACGATGTCAAGTGCCTGATCATGTTGTATGATGCCTCCCATTTGCGCATTCACGAGGAGAAGATTCTTGACAACATCAACAGTTTCACCAAGAGCTGCCTCCAGTCAGTTCTCGAAACCAACTTGGAACCGGCTCTCCAAGAAGAAGTGCGTTGCACCTTGGAGACGCCGCGTTTCCGCCGTGTTGAAAGAATCGAAGCGAAGCGCTTTATCTCCGCGTACGAAAAGAACATCGCCCGAGATGACGCCCTCTTGGAATTTGCCCGTCTGGACTACAATATCGTGCAAATTCTCTACTGCAAGGAACTGAAAGAACTTACCGTATGGTGGAAGGAGTTCCATTCCCGGACCAATCTCACCTTTGCACGAGATCGCATTGTCGAAATGTACTTCTGGGTCATGGCAATTATTTACGAGCCTTGTTACTCGTACTCCCGGATCTGGGTTACCAAAATGTTTCTATCCGTCGCCTTGTTGGACGACATCTACGACAACTATACGAGCACAGAGGAAAGCAACATCTTTACTACGGCCATGGAACGTTGGGATGTGAAGGCCACCGAACAACTCCCCGCAAACATGCGTACATTTTACGATTACTTAATTTGTACGACCGATGAGGTCGTAGAAGAATTGAAGCTTCAGAATAACAAGAATGCTGAACTTGTCAAGAAAGTCCTGATTGACGCCGCTAAATGCTACCACTCGGAGGTCAAATGGCGTGATGACCACTACGTCCCTAACGATGTTGGAGAACACCTGCAGCTTTCTATGCGTAGCATTGCCGCCATGCACTCCATCAACTTTGTCTTCATTTCTCTGGGAGCTGTGTGTACTCGTGAGGCGGTTGAATGTGCTTTCACTTACCCCAAAATTATTCGTGGTATTTGTGTTCACGCACGTATTAGTAACGACATCGCGTCACATGAGCGAGAACAGGCTTCGGAGCACATGGCCTCTACGTTGCAAACTTGCATGAAGCAGTATGGGATTACAGTCGAAGAAGCCGCTGAAAAGCTCCGTGTAATCAACGAAGAGTCCTGGATGGACATCGTTGAAGAATGCCTTTATAAGGACCAGTACCCCCTAGCGCTTTCGGAACGCGTGGTGGCTTTTGCCCAATCTATCTGTTTCATGTACAATGGTGTAGATAAATACACCATACCATCCAAACTCAAGGACAGTCTAGACTCCTTGTACGTCAACTTGATTCCTGTTATGGTCAGCAAGGGTGAGGAACTCTTTACTGGAGTCGTGCCAATCCTTGTTGAGCTTGACGGAGACGTGAACGGTCACAAGTTTTCCGTTAGCGGCGAAGGAGAAGGCGACGCCACATACGGAAAGTTGACTCTCAAGCTCATCTGCACGACGGGAAAGTTGCCGGTCCCGTGGCCCACCCTCGTCACCACCCTAGGCTACGGTTTGCAGTGTTTCGCCCGCTACCCTGACCACATGAAGCAGCATGATTTCTTTAAATCGGCCATGCCCGAAGGTTATGTCCAGGAGCGTACCATTTTCTTCAAAGACGATGGTAACTACAAGACCCGCGCTGAAGTCAAGTTTGAGGGCGACACGCTGGTAAATCGAATCGAATTGAAGGGAATCGATTTCAAGGAAGACGGCAACATTTTGGGACACAAGTTGGAATACAACTATAACTCCCACAACGTGTACATCACCGCTGACAAGCAGAAGAACGGCATCAAGGCTAATTTTAAGATCCGTCACAATATTGAAGACGGCGGTGTACAGTTGGCAGACCACTACCAACAGAACACGCCCATTGGCGACGGACCGGTCCTCTTACCGGACAACCACTATCTGTCCTACCAGAGTGCCCTCTCCAAAGACCCCAATGAGAAACGTGATCACATGGTGCTGTTGGAATTCGTTACCGCTGCAGGGATCACTCTAGGTATGGATGAACTCTATAAG

**L0_crtB-mVenus_DE 1641 bp**

AACAACCCGTCGCTTCTCAATCACGCGGTCGAGACTATGGCCGTCGGATCGAAGAGTTTTGCTACAGCCTCCAAGTTATTCGACGCAAAGACCCGGCGCAGTGTACTAATGCTCTACGCCTGGTGCCGCCATTGCGACGACGTCATTGACGATCAGACCCTCGGATTTCAGGCCAGACAGCCAGCCCTTCAAACGCCCGAACAACGTCTGATGCAACTTGAGATGAAGACCCGCCAGGCCTACGCAGGATCCCAGATGCACGAACCCGCGTTCGCCGCTTTCCAGGAAGTGGCTATGGCTCACGATATCGCCCCGGCTTACGCGTTTGACCACCTAGAAGGCTTCGCCATGGATGTACGCGAAGCTCAATACTCCCAATTGGATGATACTCTCCGATACTGCTACCACGTTGCCGGCGTTGTCGGCTTGATGATGGCCCAAATCATGGGAGTGCGTGATAACGCCACCCTCGACCGCGCCTGTGACCTTGGGCTCGCTTTTCAGTTGACCAACATTGCTCGCGACATTGTCGACGACGCGCACGCGGGCCGTTGTTATTTGCCGGCAAGCTGGCTCGAGCATGAAGGTCTTAACAAGGAGAATTACGCCGCACCTGAGAACCGTCAGGCGCTGAGCCGTATTGCCCGTCGTTTGGTGCAAGAAGCAGAACCATACTATTTGTCTGCCACAGCCGGACTGGCTGGTTTGCCCCTGCGTTCCGCCTGGGCCATCGCTACGGCGAAGCAGGTCTACCGAAAGATCGGTGTCAAAGTTGAACAGGCCGGTCAGCAAGCCTGGGATCAACGTCAGTCAACGACCACCCCCGAAAAACTGACGCTCCTCTTGGCCGCCTCTGGTCAGGCCCTTACTTCCCGGATGAGGGCTCACCCTCCCCGACCTGCCCATCTCTGGCAGCGCCCGCTCATGGTCAGCAAGGGTGAGGAACTCTTTACTGGAGTCGTGCCAATCCTTGTTGAGCTTGACGGAGACGTGAACGGTCACAAGTTTTCCGTTAGCGGCGAAGGAGAAGGCGACGCCACATACGGAAAGTTGACTCTCAAGCTCATCTGCACGACGGGAAAGTTGCCGGTCCCGTGGCCCACCCTCGTCACCACCCTAGGCTACGGTTTGCAGTGTTTCGCCCGCTACCCTGACCACATGAAGCAGCATGATTTCTTTAAATCGGCCATGCCCGAAGGTTATGTCCAGGAGCGTACCATTTTCTTCAAAGACGATGGTAACTACAAGACCCGCGCTGAAGTCAAGTTTGAGGGCGACACGCTGGTAAATCGAATCGAATTGAAGGGAATCGATTTCAAGGAAGACGGCAACATTTTGGGACACAAGTTGGAATACAACTATAACTCCCACAACGTGTACATCACCGCTGACAAGCAGAAGAACGGCATCAAGGCTAATTTTAAGATCCGTCACAATATTGAAGACGGCGGTGTACAGTTGGCAGACCACTACCAACAGAACACGCCCATTGGCGACGGACCGGTCCTCTTACCGGACAACCACTATCTGTCCTACCAGAGTGCCCTCTCCAAAGACCCCAATGAGAAACGTGATCACATGGTGCTGTTGGAATTCGTTACCGCTGCAGGGATCACTCTAGGTATGGATGAACTCTATAAG
